# Integrated single cell analysis of blood and cerebrospinal fluid leukocytes in multiple sclerosis

**DOI:** 10.1101/403527

**Authors:** David Schafflick, Chenling A. Xu, Maike Hartlehnert, Michael Cole, Tobias Lautwein, Andreas Schulte-Mecklenbeck, Jolien Wolbert, Michael Heming, Sven G. Meuth, Tanja Kuhlmann, Catharina C. Gross, Heinz Wiendl, Nir Yosef, Gerd Meyer zu Horste

**Author notes:** These authors contributed equally (David Schafflick, Chenling A. Xu, Maike Hartlehnert, Michael Cole). These authors jointly supervised the study (Nir Yosef, Gerd Meyer zu Horste). Correspondence to: Gerd Meyer zu Hörste, MD, Department of Neurology, University Hospital Münster, Albert-Schweitzer-Campus 1, Bldg A1, 48149 Münster, Germany, Nir Yosef, PhD, Department of EECS, University of California, Berkeley, 378 Stanley Hall, Berkeley 94720, USA.

## Abstract

Cerebrospinal fluid (CSF) protects the central nervous system (CNS) and analyzing CSF aids the diagnosis of CNS diseases, but our understanding of CSF leukocytes remains superficial. Here, we *firstly* provide a transcriptional map of single leukocytes in CSF compared to blood. Leukocyte composition and transcriptome were compartment-specific with CSF-enrichment of myeloid dendritic cells and a border-associated phenotype of monocytes.

We *secondly* tested how multiple sclerosis (MS) - an autoimmune disease of the CNS - affected both compartments. MS increased *transcriptional* diversity in blood, while it preferentially increased *cell type* diversity in CSF. In addition to the known expansion of B lineage cells, we identified an increase of cytotoxic-phenotype and follicular T helper (TFH) cells in the CSF. In mice, TFH cells accordingly promoted B cell infiltration into the CNS and severity of MS animal models. Immune mechanisms in MS are thus highly compartmentalized and indicate local T/B cell interaction.

## Introduction

Cerebrospinal fluid (CSF) is a clear liquid that envelops and protects the central nervous system (CNS), provides trophic support^1^ and forms a unique local immune compartment^2^. Under healthy conditions, the *non-cellular* fraction of CSF is mostly an ultra-filtrate of serum^3^ ⋅ In contrast, CSF *cells* that derive exclusively from the hematopoietic lineage exhibit a tightly controlled cellular composition considerably different from the blood^4, 5^ although the exact underlying mechanisms are unknown^6^ Leukocyte concentrations are 1,000-fold lower and CD4^+^ T lymphocytes predominate, while myeloid-lineage cells are rare compared to blood^4^ Clinically, CSF provides a unique diagnostic window into inflammatory and degenerative diseases of the CNS, but its low cell concentration and limited available sampling volume have thus far impeded transcriptional analyses of CSF cells and a comprehensive characterization of single CSF cells in inflammatory CNS diseases is unavailable^7,8^.

Single-cell transcriptomics is a transformative and rapidly evolving technology that has mostly been used to re-define the heterogeneity of complex tissues from healthy rodents or humans^9–11^ ⋅ Diseased tissues have also been analyzed with single-cell technologies and the cancer field has seen their rapid adaptation^12, 13^. Proponents of the technology posit that insights from single-cell transcriptomics are likely to translate into palpable benefits for human patients and enable precision medicine in the not-too-distant future^14, 15^. However, outside of the field of cancer, just a handful of studies have used the technology to compare tissue samples from disease-affected vs. control donors in a clinically relevant setting^16^. This leaves many methodological and conceptual questions unexplored.

Here, we applied single-cell transcriptomics to blood and CSF cells from patients with multiple sclerosis (MS) and controls, validating key findings with flow cytometry and mouse model studies. MS is a chronic inflammatory, demyelinating disorder of the central nervous system (CNS)- most likely of autoimmune origin - causing substantial disability ^17^ We chose this paradigmatic inflammatory disease, because many questions remain unanswered despite a vast amount of available literature. In fact, evidence supports the involvement of both T cells and B cells in MS, but the relative contribution of each cell type to disease aetiology is unknown. On the one hand, production of immunoglobulins and expansion of B lineage cells^4, 18^ occurs in the CSF with evidence of antigen-driven maturation^19, 20^ and B cell depleting therapies are effective in MS^21^. On the other hand, T cells are abundant in MS lesions^22, 23^ and T cells are affected by many established MS treatments and induce an MS-like condition named *experimental autoimmune encephalomyelitis* (EAE) in rodents^24^. Therefore, much is to be learned about the relative contribution and interaction of T cells with B cells in MS especially in the CSF.

We speculated that single-cell transcriptomics would help fill this knowledge gap by identifying a compartment-specific and disease-specific composition and transcriptome of rare CSF cells. In fact, we here describe a previously unknown enrichment of myeloid dendritic cells and regulatory T cells in the CSF, while other lineages including NK cells and B cells are less abundant. In MS, we find and independently confirm an expansion of NK cells and late-stage B lineage cells that is restricted to the CSF. We also introduce a new analytical approach termed cell set enrichment analysis (CSEA) to identify cluster-independent transcriptional findings and thereby observe an expansion of B cell-helping T follicular helper (TFH) cells. In a reverse-translational approach, we confirm that such TFH cells promote CNS autoimmunity and local B cell infiltration in two distinct animal models of MS. We thus demonstrate how an unbiased approach can aid our understanding of a unique human immune compartment and identify cellular mechanisms locally driving CNS disease.

## Results

### Single cell transcriptomics reconstructs cell types in cerebrospinal fluid and blood

We first aimed to identify the compartment-specific composition and expression of CSF cells compared to blood using an unbiased approach (Fig. 1A). We recruited patients with idiopathic intracranial hypertension (IIH) as controls and treatment-naïve patients with clinically isolated syndrome (CIS) or relapsing-remitting MS (together termed MS, Methods) donating blood and CSF. Both cohorts were well matched and CSF parameters were either comparable between groups or exhibited known MS-associated changes (Suppl. Fig. 1A-D, Suppl. Tab. 1-2). Using microfluidics-based single cell RNA-sequencing (scRNA-seq) we obtained in total 42,969 blood single cell transcriptomes (5 control vs. 5 MS donors) and 22,357 corresponding CSF single cell transcriptomes (4 control vs. 4 MS donors). Genes detected per donor were 934.4 ± 379.1 SEM in PBMCs and 1,021.4 ± 374.0 SEM in CSF (Suppl. Tab. 3). After filtering and normalization, we performed multi-step clustering of the merged 65,326 blood/CSF cell dataset (Suppl. Fig. 2A). We thereby classified 61,051 single cells into 17 final cell clusters (Fig. 1B, Suppl. Fig. 2A). Based on marker gene expression (Fig. 1C, Suppl. Fig. 2D, Suppl. Tab. 4; selected protein names in non-italic), we identified αβ T cells *(CD3E, LCK, TRAC, TRAJ16)* subsetting into CD4+ T cells *(IL7R, CD4)*, activated CD8+ T cells (*COBB, CCL5)*, naive CD8+ T cells (*CD8B*, CCR7), regulatory T cells *(FOXP3, CTLA4)* and a small cluster of γδ T cells *(TRDC).* Two NK cell clusters *(GNLY, NKG7)* most likely represented the more cytotoxic and mature CD56^dim^ (NK1; FCGR3A/CD16, *PRF1)* and more naive CD56^bright^ (NK2; *SELLICD62L, XCL1)* subsets. Three B lineage clusters *(CD74, CD79A, IGH* gene family) corresponded to naive B cells (B1; *CD37, IGHO)*, activated B cells (B2; *CD27, IGHM)*, and plasma blasts (plasma; *IGHG, CO3B, TNFRSF17/CD269;* negative for *MS4A1/CD20*, SOC1/CD138). Myeloid lineage cells *(LYZ)* separated into myeloid dendritic cells (mDC) type 1 (mDC1; *WOFY4, XCR1, BATF3)*, mDC type 2 (mDC2; *FCER1A, CD1C, CLEC10A)*, and granulocytes (granulo; *S100AB, S100A9).* Using canonical markers, the two identified monocyte clusters represented classical (ncMono; *CD14)* and non-classical (Mono; *FCGR3AICD16)* monocytes although the two clusters were mostly separated by tissue of origin (see below). Additional clusters represented plasmacytoid dendritic cells (pDC; *TCF4/E2-2, TNFRSF21/DR6)* and megakaryocytes (MegaK; *GNG11, CLU).* Microfluidics-based scRNA-seq thus successfully reconstructs leukocyte lineages from CSF and blood.

**Figure 1:**
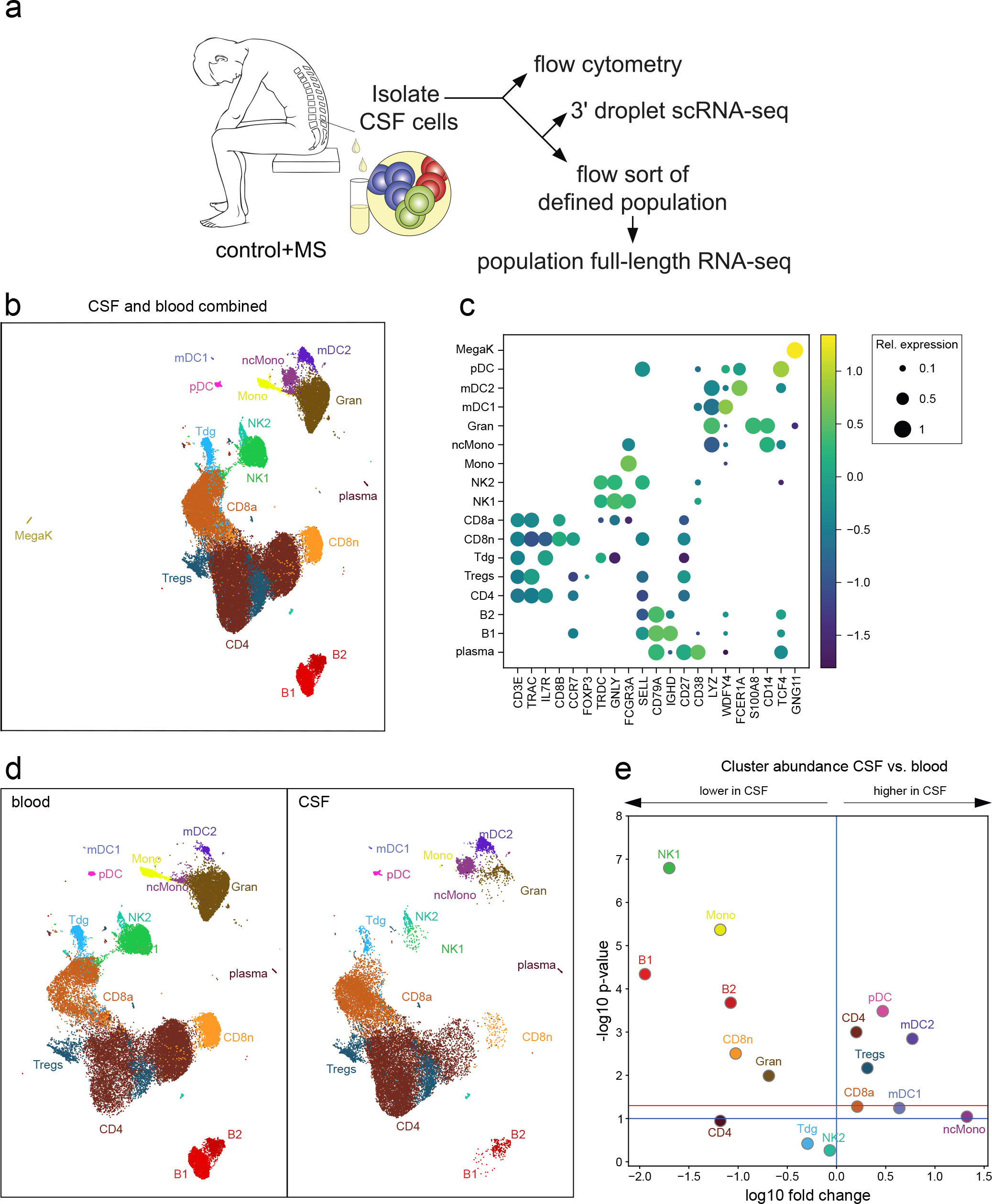
Single-cell transcriptomics reconstructs the compartment-specific leukocyte composition of CSF and blood. (A) Schematic of the study design (Methods). (B) Uniform Manifold Approximation and Projection (UMAP) plot representing 17 color-coded cell clusters identified in merged single cell transcriptomes of blood (42,969) and CSF (22,357) cells from control (n = 4) and multiple sclerosis (MS) (n = 4) patients (Methods). Cluster names were manually assigned. (C) Dotplot depicting selected marker genes in cell clusters. Dot size encodes percentage of cells expressing the gene, color encodes the average per cell gene expression level. (D) UMAP plots comparing blood (left) and CSF (right) cell clustering. Please note that the MegaK cluster is disregarded for higher resolution. (E) Volcano plot depicting differences of cluster abundance in CSF compared to blood plotting fold change (log10) against p-value (-log10) based on beta-binomial regression (Methods). Horizontal line indicates significance threshold. Cluster key: pDC plasmacytoid dendritic cells (DC), mDC1 myeloid DC type 1, mDC2 myeloid DC type 2, ncMono non-classical monocytes, mono monocytes, gran granulocytes, Tdg γδ T cells, CD8n naive CD8^+^ T cells, CD8a activated CD8^+^ T cells, Tregs regulatory CD4^+^ T cells, CD4 CD4^+^ T cells, NK natural killer cells, MegaK megakaryocytes, B1 / B2 B cell subsets, plasma plasmablasts.

### Cerebrospinal fluid leukocytes exhibit a compartment-specific composition and transcriptome

CSF cells have not been characterized with unbiased approaches. We therefore next analyzed the compartment-specific cell type composition identified by unbiased scRNA-seq in CSF compared to blood. As expected for CSF^4, 25^, non-hematopoietic cells (e.g. neurons, glia, ependymal cells), megakaryocytes, granulocytes, and RBC (removed from final clustering) were absent or strongly reduced compared to blood (Fig. 1D,E, Suppl. Fig. 3A,B). We also found CD56^dim^ NK1 cells reduced among CSF cells, while the NK2 cluster was not different (Fig. 1D,E). Both the mDC1 and mDC2 clusters had a significantly higher proportion in CSF than in blood (Fig. 1D,E). Notably, mDC1 cells expressed markers indicating cross-presenting capacity *(XCR1, WOFY4* ^26^; Fig. 1C). Among T cells, total CD4 cells and Tregs were more abundant in the CSF, while CD8 T cell clusters were not different (Fig 1D,E). Flow cytometry confirmed this unique composition of CSF leukocytes (Suppl. Fig. 4A-C. Cell proportions in CSF and blood did not correlate by either scRNA-seq or flow cytometry (data not shown) supporting an independent regulation of their cell composition. In summary, we confirm a highly compartment-specific composition of CSF cells and identify a previously unknown enrichment of mDC1 and Tregs in the CSF.

We also found a prominent CSF-specific pattern of myeloid lineage cells. The cluster tentatively named ncMono was almost exclusively CSF-derived, did not co-cluster with blood monocytes (Fig. 1D, Suppl. Fig 2C), and was transcriptionally distinct (Suppl. Tab. 4). Expression of canonical markers indicated a CD14^+^FCGR3A/CD16^int^ phenotype of the ncMono cluster (Fig. 1C) as described for CSF monocytes ^27^. Additionally, we identified a unique transcriptional signature that was not captured by the dichotomous classical/non-classical classification. The CSF-derived ncMono cluster expressed genes previously identified in classical (CD9, *CD163, EGR1, BTG2*) and in non-classical *(C1QA, C1QB, MAF*, CSF1R/CD115) monocytes^28^⋅ Notably, the ncMono cluster also expressed (Suppl. Tab. 4) markers of perivascular macrophages *(LYVE1;* ^29^), microglia (*TREM2, TMEM119, GPR34;* ^30^) and CNS border associated macrophages *(STAB1, CH25H;* ^31, 32^) previously identified in rodents. We thus identify a distinct phenotype of CSF monocytes.

We next aimed to identify further compartment-specific gene expression signatures on a per cluster level (Suppl. Tab 5). We focussed on genes identified independently as differentially expressed (DE) by two methods (Mann-Whitney U test, edgeR ^33^) *and* supported by Bayesian model comparison in scVI (34, Methods). Due to the stringency of this approach, most of such ‘triple-consistent’ genes were DE in CSF vs. blood cells in only one (18.9% of all expressed genes) or two (5.1%) clusters (Suppl. Tab 5), although measures of differential expression were positively correlated especially between related clusters (Suppl. Fig. 10A) indicating coregulated gene modules in related cell types.

Genes induced in multiple (i.e. >3) CSF clusters included FGF9, previously implicated in inflammatory CNS tissue damage^35^ and Metallothionein E, potentially involved in CSF metal ion homeostasis^36^. Cell cycle (e.g. CCNC/Cyclin-C) genes were induced in CD4^+^ T cells in line with their activated phenotype in CSF^37, 38^. Genes induced in CD4^+^ T cells in the CSF were also related to lipid antigen recognition (*CD1E)*, interaction with antigen-presenting cells *(CD81, CD83, CD84, CD209)* and adhesion and migration (CD99). In fact, CSF T cells expressed a specific pattern of chemokine and integrin transcripts including an induction of *CXCL16* and *CXCR5* and downregulation of ITGALNLA4 in CSF CD4^+^ T cells and of ITGB7 in myeloid cells (Suppl. Fig. 3C, Suppl. Tab. 5). Genes consistently downregulated in CSF T were associated with naïive cell state (SELL/CD62L), cytokine responses *(/L2RG/common* y chain). Interestingly, CD48 previously associated with CSF translocation of bacteria^39^ was upregulated in CSF T cells (Suppl. Tab. 5). In accordance, GSEA showed enrichment of pathogen response pathways in CSF induced genes (e.g. KEGG pathways hsa05169, hsa05168) (Suppl. Tab. 6). Single cell transcriptomics thus identifies a unique phenotype of CSF leukocytes and specific pattern of trafficking molecule expression.

### Multiple sclerosis preferentially alters transcription of blood and composition of CSF cells

Next, we analysed our dataset for MS-associated changes. Blood cells exhibited no significant differences in composition in MS compared to control (Suppl. Fig. 5A,B) as confirmed by flow cytometry (Suppl. Fig. 4B,C). In contrast, blood cells exhibited diverse ‘triple-consistent’ (see above and Methods) transcriptional changes (Suppl. Tab. 7) including an induction of activation markers (/COS), specific cytokine receptors *(IL17RA)*, and trafficking molecules *(PECAM1/CD31, ITGA5/α5* integrin) in T cells (Suppl. Fig. 5C).

In contrast to blood that lacked compositional changes, the cell type composition of CSF was clearly different in MS patients compared to controls (Fig. 2A,B). Using binomial regression modelling (Methods), all B lineage cell clusters (B1, B2, plasma) significantly expanded in the CSF in MS compared to controls (Fig. 2A,B) in accordance with flow cytometry (Suppl. Fig. 4B,C) and previous studies^18,40,41^. Heavy chain gene expression in mature B cell clusters (B2, plasma) was dominated by IGHG/lgG genes, although some cells expressed IGHA/lgA genes (Suppl. Fig. 6A-D). The proportion of cells expressing the K- and λ-light chain was at an average IGKC/κ-to-lGLC/λ ratio of 2.75 in CSF compared to 1.92 in blood in line with expectations. Most B lineage cells in the CSF are thus class-switched because heavy chain usage in blood evolves from *IGHD* to *IGHM* to *IGHG/IGHA* during activation and maturation. Among other cell lineages, both CD56^dim^ NK1 and CD56bri NK2 cell clusters and naive CD8 cells increased in the CSF in MS compared to controls (Fig. 2A,B) as confirmed by flow cytometry (Suppl. Fig. 4B,C) and in line with a previous study ^42^ In addition, we identified a previously undescribed increase of mDC1 cells and Tregs in the CSF in MS, while yoT cells (Tdg) were significantly decreased (Fig. 2A,B). Alternative t-test statistics returned comparable results (data not shown). MS thus induces complex changes of the composition of CSF leukocytes that are characterized by a simultaneous expansion of cell types with the capacity for antibody production (B1, B2, plasma), cytotoxicity (nCD8, CD56^dim^ NK1) and with regulatory potential (Tregs, CD56^bri^ NK2).

We next tested for disease-associated ‘triple-consistent’ transcriptional changes in CSF cell clusters (Suppl. Tab. 8). In CSF T cells, we found an induction of genes associated with immune activation *(HLA-C*, CD5) and with interferon responses *(/L12RB1, IL18RAP)* and related down-stream signaling molecules *(IRF3, IRFB)* (Fig. 2C). Specific trafficking molecules (e.g. ITGB1/integrin-β1) were also up-regulated in MS. The Treg cluster showed induction of the transcription factor *STAT1* and some interferon-regulated genes *(MUM1, NUCB2).* The mDC2 cluster induced B cell related genes (e.g. *CD79A, CD74)* and signs of IL-2 signaling *(STAT5A)* and a co-inhibitory molecule *(TNFRSF18/GITR).* The pDC cluster expressed more *IRF7* - upstream regulator of interferon signaling. The NK2 cluster showed induction of hypoxia-sensing transcripts *(HIF1A).* The MS-associated cellular response in CSF is thus diverse and lineage specific and shows signs of interferon-regulated responses (Suppl. Tab. 8).

**Figure 2:**
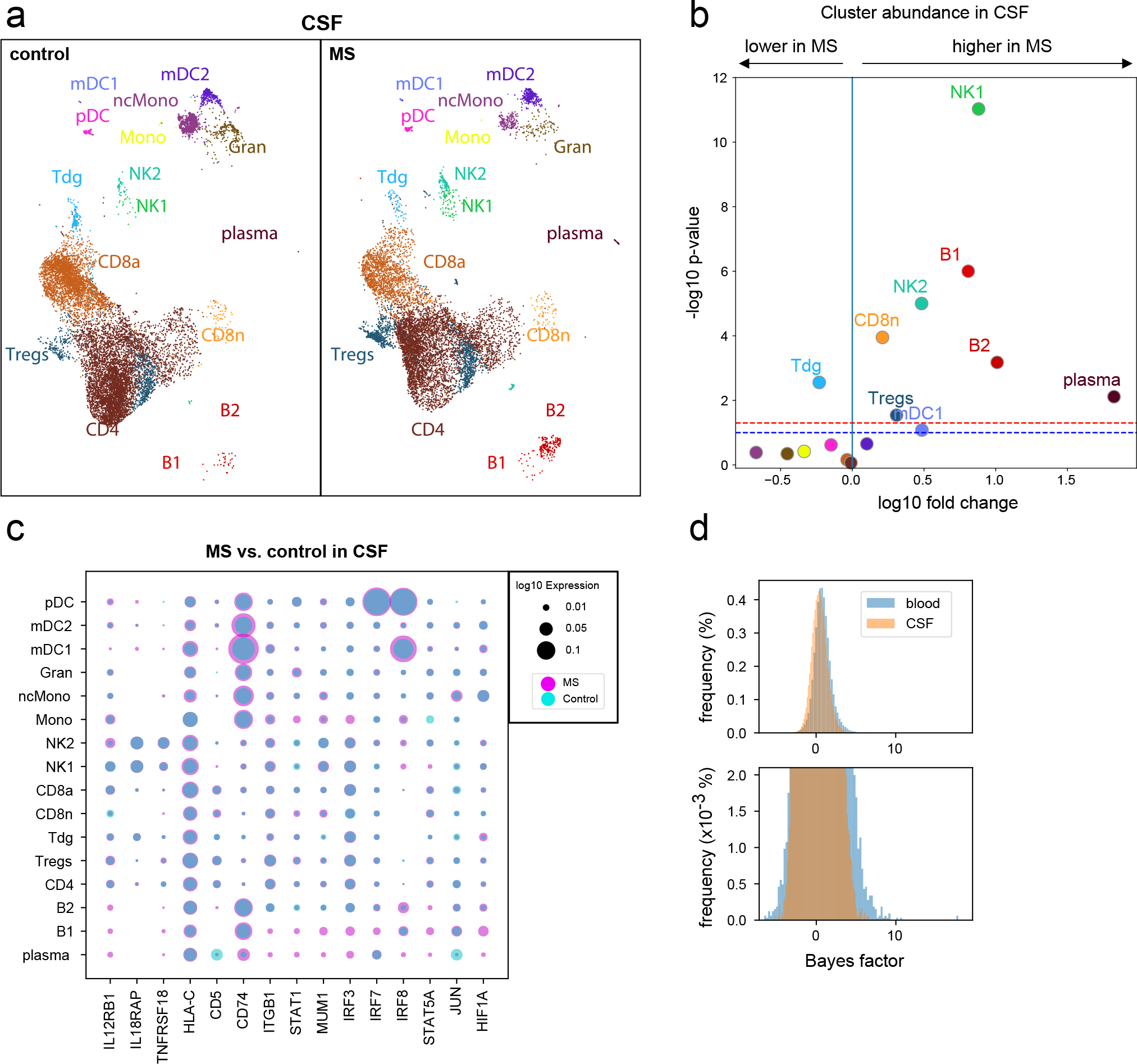
MS predominantly alters CSF cell composition and blood cell transcription. (A) Comparative UMAP plots depicting only CSF cells from control (12,705 cells, left plot) and MS (9,652 cells, right plot) donors. Color coding and cluster names are as in Figure 1. (B) Volcano plot showing differences of cluster abundance of only CSF cells in MS samples compared to controls plotted as fold change (log10) against p-value (-log10) based on beta-binomial regression. (C) Dotplot depicting selected genes differentially expressed in at least one cluster of MS cells compared to controls in CSF. Dot size encodes percentage of cells expressing the gene. Purple indicates higher, turquoise indicates lower expression in MS, respectively. (D) Bayes Factor (BF) frequency histogram in all cluster-specific case-control differential expression (DE) analyses colored by tissue. Higher magnification in bottom panel. Only clusters with a minimum of 10 cells per tissue per disease state are included. Please note that the BF is proportional to the likelihood of differential expression (i.e. higher BF indicates more likely DE)^34^.

When directly comparing disease effects between CSF and blood, we found that - surprisingly - a considerably greater proportion of expressed genes was differentially expressed in the MS condition in blood than in CSF. For example, 354 genes were DE in MS compared to control within the blood CD8a cell cluster vs. 24 genes in the same cluster in the CSF (Suppl. Tab. 7,8). Overall, when plotted across all clusters and genes, the Bayes factor (a measure of likelihood of differential expression that does not depend on sample size) of the MS vs. control comparison showed more extreme values in blood than in CSF (Fig. 2D). Then we subsampled each cluster to have the same number of cells in blood and CSF and ran the Mann-Whitney U test and observed that the blood case-control had more significant P-values and those P-values were more extreme (data not shown). In blood, MS thus preferentially increases *transcriptional* diversity, while in CSF it preferentially increases *cell type* diversity suggesting compartment-specific disease mechanisms.

### T helper cells with cytotoxic phenotype are increased in multiple sclerosis

We had tentatively handled the CD4^+^ T cell cluster as one cell type, because this population did not form clearly distinct sub-clusters (Suppl. Fig. 2A) and because many well-established T cell protein markers faired poorly on transcript level. We therefore next aimed to better characterize the CD4^+^ T cells using dedicated approaches.

We performed sub-clustering of the CD4^+^ T cell cluster (Fig. 3A, Suppl. Fig. 7A-C, Suppl. Tab. 9). As expected for an unsupervised clustering approach^43^, we found a minor population of CD8 T cells *(CD8B;* CD4^+^ T cell sub-cluster (CD4Tc) #8; 7.54% of all CD4^+^ T cells) ‘remaining’ within the tentative CD4^+^ T cell cluster (Fig. 3A,B). The CD4^+^ T cells broadly separated into naïve-like (*SELL, CCR7;* CD4Tc #5,11,1,2) and memory-like (*CO44;* CD4Tc #9,4,0,3,6,7) clusters based on marker gene expression (Fig. 3B). Memory cells further separated into subsets with mostly effector memory-like *(CD69;* CD4Tc #3,0,4) and central memory-like (CD27; CD4Tc #7,6,9) phenotype. We also identified a cluster of likely Treg identity *(FOXP3, CTLA4*, CD4Tc #10, Fig. 3B) located at the intersect between naïve and memory cells (Fig. 3A). Notably, this cluster expressed individual markers of T cell exhaustion *(TIGIT)* (Suppl. Fig. 7F)^44^ previously associated with loss of suppressive capacity of Tregs in the tumour micro-environment^45^.

**Figure 3:**
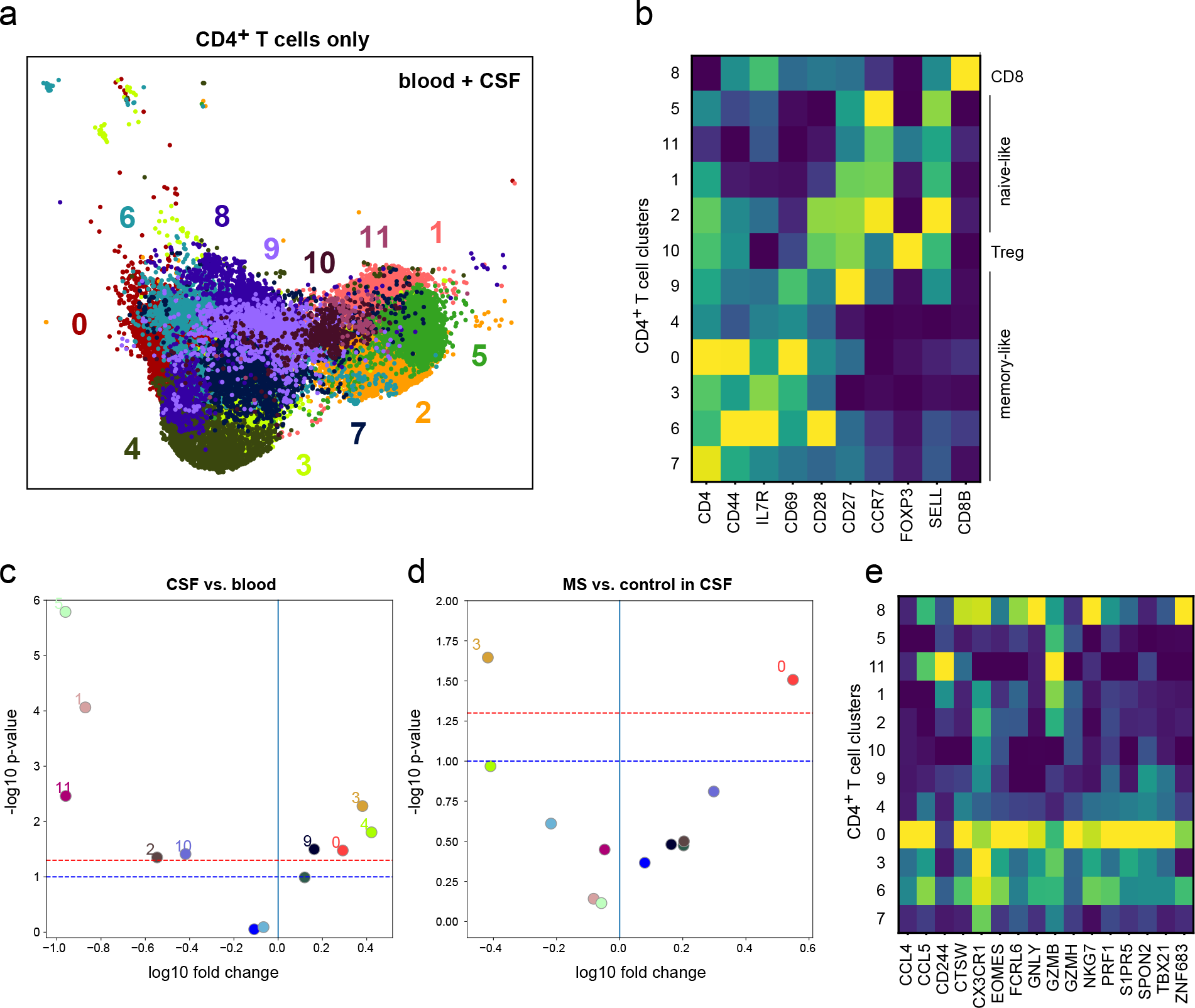
Cytotoxic-like population of CD4^+^ T cells is induced in the CSF in MS. (A) UMAP plot showing sub-clustering of all CD4^+^ T cells combined from blood (13,933 cells) and CSF (11,172 cells). Sub-clusters are numbered 0-11. (B) Heatmap depicting per cluster average expression of selected T cell subset marker genes. (C) Volcano plot showing differences of CD4^+^ T cell cluster abundance in CSF compared to blood as fold change (log10) against p-value (-log10) based on Student’s t-test. (D) Volcano plot showing differences of CD4^+^ T cell cluster abundance in MS compared to control within CSF based on Student’s t-test. (E) Heatmap showing average gene expression of selected cytotoxicity markers derived from ^48^.

We next used VISION (previously named FastProject^46^) to identify transcriptional signatures rather than individual marker genes to better interpret the CD4^+^ T cell sub-clustering. Transcriptional signatures identified a transcriptional gradient ranging from naive to memory T cell state (Suppl. Fig. 7D). This is in line with previous findings in rodents ^47^, and may indicate that CD4^+^ T cells generally form transcriptional gradients rather than distinct subclusters also explaining the poor applicability of clustering approaches alone for this cell type.

We next sought to identify compartment- and disease-specific changes among CD4^+^ T cell sub-clusters. We found that several memory-type clusters (CD4Tc #3,4,0,9) were more abundant in CSF compared to blood while naïve clusters (CD4Tc #1,11,2) and exhausted Tregs (CD4Tc #10) were less frequent using t-test based statistics (Fig. 3C; Suppl. Fig. 7E) in accordance with previous studies^37, 38^. Disease-associated changes in blood were limited to a reduction of a single memory-like cluster (CD4Tc #4) in MS compared to control (Suppl. Fig. 7E). Transcriptional changes in blood and CSF did not encompass any of the key T helper cell lineage transcripts (e.g. *TBX21, GATA3, RORC*, Suppl. Tab. 10,11). In CSF, a CD4^+^ T cell sub-cluster (2,240 cells) of memory cells was significantly more abundant in MS vs. control (CD4Tc #0; Fig. 3D). This cluster expressed multiple genes associated with cytotoxic function *(GZMB, PRF1, CCL5)* despite similar levels of CD4^+^ T cell marker genes *(CD4, IL7R)*, low doublet probability (predicted doublet t-test p value 0.68), and absence of CD8 or NK cell markers *(COBB, NKG7;* Fig. 3E) in this population. This gene signature showed considerable similarity with a recently described population of cytotoxic CD4^+^ T cells^48^ that is enriched within the CD4^+^ T cells effector memory recently activated (TEMRA) compartment. This indicates that cytotoxic CD4^+^ T cells expand in the CSF in MS.

### Cell set enrichment analysis (CSEA) identifies cluster-independent transcriptional changes

Although the clustering analysis was informative about the general other states, it was not readily able to identify a stratification of the cells into specific T helper cells subsets. Indeed, using a transcriptome-wide measure of similarity to stratify the cells may be insensitive to subtle differences in cell state. This may result in cells from the same state not being assigned to the same cluster. Another possible scenario is that cells from the same state will be primarily assigned to the same cluster, but with many additional cells. Both of these cases may lead to missing functional properties that are specific to MS, particularly in cases where these properties are present in a small subset of cells. We therefore developed a novel procedure – cell set enrichment analysis (CSEA) – which reutilizes the GSEA test for working on ranked lists of cells rather than genes (Methods, Suppl. Fig. 7). In this procedure, the cells are first ordered by a transcriptional phenotype of interest (e.g., summed expression of genes in a pathway). The statistical test can then detect cases in which a subset of cells from one group (e.g., MS) exhibit unusually high or low values of that transcriptional phenotype compared to cells from the second group (e.g., control).

We used this analysis with signature scores obtained from the VISION pipeline based on signatures obtained from databases and literature curation (Methods) to specifically analyze CD4^+^ T cells from CSF and blood.

Our CSEA testing procedure returned lists of cell sets significantly (Methods) enriched in MS and expressing a certain gene signature (Suppl. Tab. 12). The cell sets that were enriched in MS when compared to controls expressed signatures of T helper cell type 1 (Th1)^49^ and T follicular helper (TFH) cells^50^ (Suppl. Fig. 8B,C). We validated the results by testing against random genesets that are specifically matched to the signature sets (Methods) and found that the TFH signature was enriched in the CSF (*P* = 0.002) but not in the blood (*P* = 0.889). Th1 cells are significantly enriched in both blood (P=0.012) and CSF (p=0.0). The leading edge size reflects the number of cells driving the high enrichment score (ES). In all cases the leading edge is small (<600 cells) (Suppl. Tab. 12) indicating that a subset of cells is driving the enrichment. Thus, CD4^+^ T cells expressing a Th1- and TFH-like signature are enriched in MS in the CSF, but are spread across sub-clusters. Our novel analytical approach can therefore decouple clustering of cells from disease-state or differentiation-state enrichment of cells, providing a new framework for interpreting complex scRNA-seq datasets. Interestingly, TFH cells are required for B cell maturation ^51^. This lead us to hypothesize that TFH might be functionally related with the MS-specific B cell-expansion in the CSF.

### B cell-helping T follicular helper cells expand in the CSF in multiple sclerosis and exacerbate corresponding animal models

We therefore next tested whether TFH cells are in fact altered in the CSF in MS. We identified CD3^+^CD4^+^CXCR5^+^ TFH cells in the CSF by flow cytometry and found a significantly increased proportion of TFH cells in MS patients (Fig. 4A,B) in accordance with previous studies in the blood^52, 53^ and CSF^54^. Activated PD-1^+^ and PD-1^+^ICOS^+^ TFH cells were also increased in the CSF (Fig. 4B) while the alternative CD4^+^CXCR5^−^PD-1^+^ subset^55^ was unchanged (data not shown). The percentage of PD-1^+^ TFH cells in CSF positively correlated with the proportion of CSF plasma cells quantified by flow cytometry (r = 0.70, p < 0.05). Next, we performed bulk population RNA-seq from sorted TFH cells from the CSF of MS patients (n = 9) vs. controls (n = 9) to better characterize this cell type. Surprisingly, no genes reached the significance threshold for differential expression (Suppl. Tab. 13). This indicates that CSF-resident TFH cells increase in abundance in MS, but do not alter their phenotype.

**Figure 4:**
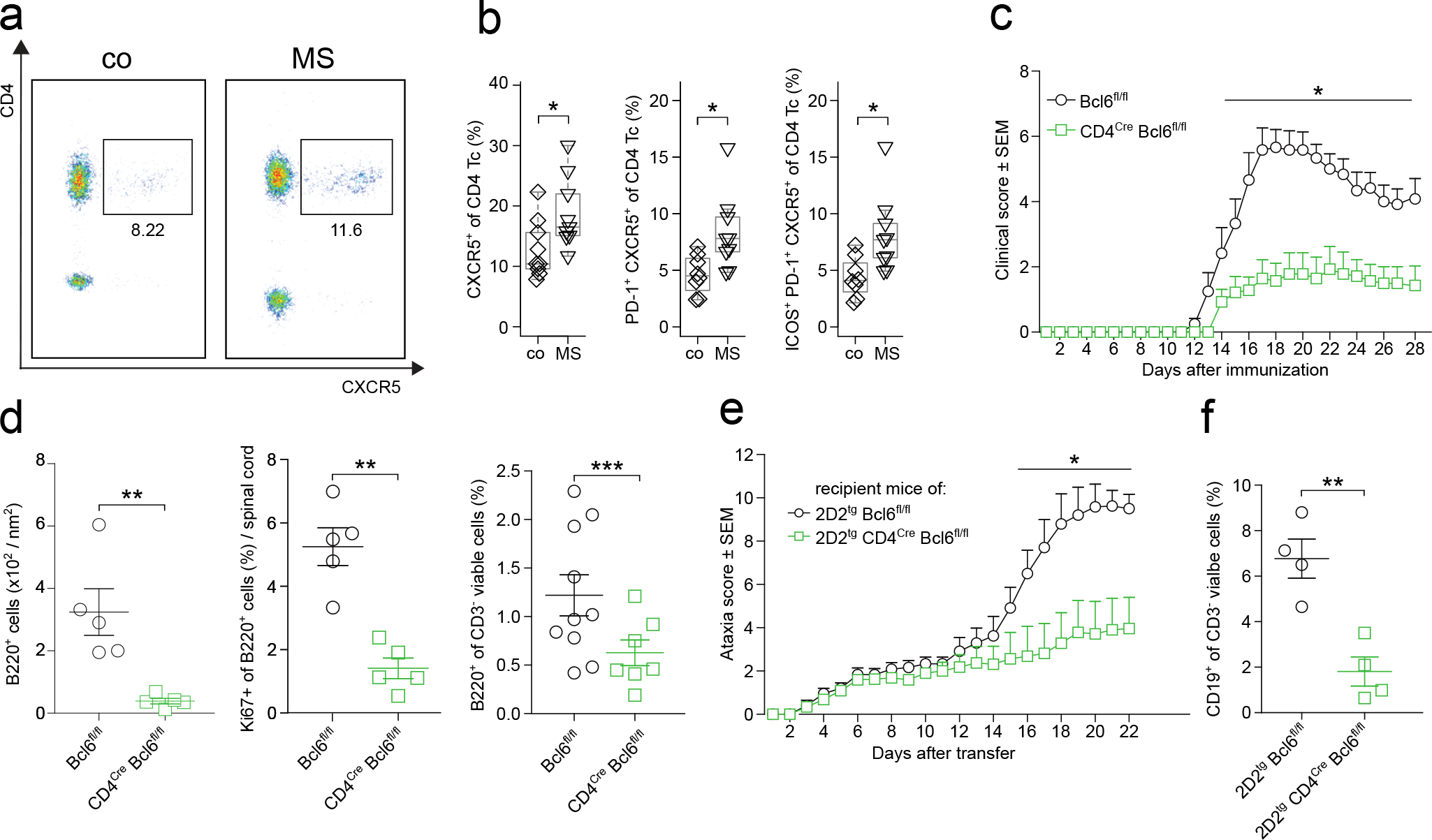
T follicular helper (TFH) cells expand in the CSF in MS and promote MS animal models. (A) Representative flow cytometry dot plot of CSF cells from a control and MS patient after gating on live CD3^+^ cells. (B) Proportion of CXCR5^+^ (left), of PD-1^+^CXCR5^+^ (middle), and of ICOS^+^PD-1^+^CXCR5^+^ (right) cells among live CD3^+^CD4^+^ T cells in CSF of control (co; n = 9) and MS (n = 9) patients quantified by flow cytometry. (C) Active EAE was induced in Bcl6^fl/fl^ (wildtype, circles, n = 6) and CD4^Cre^ Bcl6^fl/fl^ (squares, n = 7) mice using MOG_35___55_ peptide (Methods). Mice were monitored daily for clinical EAE signs. One representative of three independent experiments is shown. (D) At day 28 after EAE induction, the density of CD3^−^B220^+^ leukocytes was quantified in spinal cord paraffin cross-sections by histology (left). The proportion of Ki67^+^ among B220+ cells was quantified (middle). The proportion of B220+ cells was quantified by flow cytometry at peak of EAE (right). (E) Naive CD4^+^ T cells were sorted from Bcl6m^fl/fl^ 2D2^tg^ mice (circle) and CD4^Cre^ Bcl6^fl/fl^2D2^tg^ mice (squares), differentiated *in vitro* (Methods), and intravenously injected into wildtype recipient mice (n = 6-8 per group) at 5×10^6^ cells per mouse. Recipients were monitored for signs of EAE. One representative out of three independent experiments is shown. (F) At day 28 after transfer, the proportion of CD3^−^CD19^+^ leukocytes in brain and spinal cord was quantified by flow cytometry. * *P* < 0.05, ** *P* < 0.01, *** P < 0.005.

We then tested whether TFH cells in fact promote other-inflammation to a functionally relevant extend using common animal models of MS. We generated mice with T cell-restricted deficiency of Bcl6 - the lineage-defining transcription factor of TFH cells^51^. Such CD4^*Cre*^Bcl6^*fl/fl*^ mice lack TFH cells and fail to mount antigen-specific B cell responses^51^, while the differentiation of other T helper cell lineages (Suppl. Fig. 9A) and the composition of the peripheral immune compartment after immunization were unchanged (Suppl. Fig. 9B) as previously described^56^.

We induced active EAE using myelin oligodendrocyte glycoprotein (MOG)_35-55_ peptide in these mice and EAE severity was significantly reduced in CD4^*Cre*^Bcl6^*fl/fl*^ mice compared to ere-negative littermates (Fig. 4C). Accordingly, the number of inflammatory lesions and infiltrated area in the spinal cord of CD4^*Cre*^Bcl6^*fl/fl*^ mice were lower than in controls (Suppl. Fig. 9C,D). We tested how the absence of TFH cells influenced B cells in the CNS and found a lower proportion of B cells (B220^+^ CD3^−^) infiltrating the CNS in CD4^*Cre*^Bcl6^*fl/fl*^ mice by flow cytometry (Fig. 4D) and in the spinal cord by histology (Fig. 4D, Suppl. Fig. 9E). To reliably exclude a relevant contribution of Bcl6-deficiency on the priming phase of EAE, we next generated 2D2^*tg*^ CD4^*Cre*^Bcl6^*fl/fl*^, mice expressing a T cell receptor transgene recognizing MOG^57^ to enable immunization-independent adoptive transfer EAE induction. After transfer of interleukin (IL)-17 producing myelin-reactive T helper cells into wildtype hosts (Methods), 2D2^*tg*^ Bcl6^*fl/fl*^ control T cells induced considerably more severe EAE than Bcl6-deficient 2D2^*tg*^ CD4^*Cre*^Bcl6^*fl/fl*^ donor cells (Fig. 4E). Control recipients also showed a higher proportion of B cells in the CNS than recipients of Bcl6-deficient T cells (Fig. 4F). Taken together, our data indicate that TFH cells locally drive B cell responses in the CNS and promote MS-like autoimmunity.

## Discussion

In this study, we constructed the first unbiased comparative single-cell map of blood and CSF cells. We identify a compartment-specific leukocyte transcriptome and composition including a previously unknown enrichment of mDC1 and Tregs in the CSF. Monocytes in the CSF were especially distinct and partly resembled CNS border-associated macrophages. These findings emphasize the unique immune microenvironment of the CSF. We used MS to test how a paradigmatic autoimmune disease would affect leukocytes in a compartment-specific fashion. Surprisingly, we found that MS preferentially increased *transcriptional* diversity in blood, while it increased *cell type* diversity in CSF thus providing evidence for compartmentalized mechanisms driving human autoimmunity in the brain. In MS-derived CSF we found an expansion of cells resembling a recently described subset of cytotoxic CD4^+^ T cells^48^. We also found that clustering-based methods alone poorly capture disease-associated changes within CD4^+^ T cells and developed CSEA as a new cluster-independent analytical approach to identify TFH cells increased within the CSF. Such TFH cells in fact promoted local B cell accumulation and disease severity in MS-like animal models. Our study thereby also provides a signature case for reverse translation from unbiased transcriptomics in humans to disease mechanisms in rodents.

Our unbiased approach considerably extends the available flow cytometry-based characterization of CSF leukocytes^4, 25^ and is the first to identify local enrichment of rare cell types, such as Tregs and the mDC1 subset. Notably, mDC1 cells abundant in the CSF expressed markers of cross-presenting capacity *(XCR1, WDFY4;* ^26^) while NK2 cells in the CSF expressed the corresponding ligands *(XCL1, XCL2)* indicating that cell types equipped for cross-presentation and anti-viral defence circulate the CSF. We also replicate the known activated/memory phenotype^6, 38^ of CSF-resident T cells and identify a distinct pattern of adhesion molecule expression in CSF leukocytes (Fig. 3E). We also generated a repository of compartment-specific gene expression signatures for potentially specifically targeting CSF cells in the future (e.g. CCL3 in CSF myloid cells (Suppl. Fig. 3C)). This also allowed us to provide a human confirmation beyond a previous single case study in HIV^8^ of the rodent border-associated macrophage cell phenotype^31, 32^. Our findings thus lend further support to a species-independent ‘peri-CNS immune system’ involved in local autoimmunity and anti-pathogen defense.

A plethora of studies have analyzed mechanisms of neuro-inflammation^58, 59^ albeit often equating rodent models with human MS. Unlike our study, purely human studies often solely relied on easily accessible peripheral blood mononuclear cells^60^ sometimes even using unsorted cells^61^. Some transcriptional studies of blood cells focussed on T cells^62^, different treatments^63−65^, or myelin antigen-specific T cells using pre-defined gene-sets^60^. However, whether blood leukocytes actually constitute a suitable surrogate of disease mechanisms in MS remains unknown. A single available transcriptomic study of unsorted bulk CSF cells in MS returned signs of local B cell expansion^7^ and a comprehensive transcriptional profiling in MS at sufficient resolution is lacking. In generating such a characterization, we here surprisingly observe that MS differentially affects the blood vs. CSF immune compartments. It appears that MS increases *transcriptional* entropy in blood, but *cellular* entropy in CSF potentially indicating compartment-specific disease mechanisms. It remains to be tested whether similar compartment-specificity also occurs in other human autoimmune diseases.

Specific T helper (Th) cell lineages have long been other with MS-like pathology in rodents, while evidence in human MS is more ambiguous^66, 67^. Notably, *blood* T cells showed some induction of Th17 cell-related signalling *(/L6R)* although most core Th17 transcriptional modules were not differentially expressed^68^. In contrast, CSF cells showed signs of Th1 cell-related signalling on the individual gene level (e.g. *IL12RB1, IL18RAP, IRFB)*, by GSEA (Suppl. Tab. 6), and when using CSEA (Suppl. Fig. 8B). In addition, we found a previously undescribed expansion of CD4^+^ T cells with cytotoxic phenotype in the CSF in MS (Fig. 3E). We thus speculate that cytotoxic CD4^+^ T cells may be involved in local MS pathology in the CSF. We also found that in accordance with previous studies^47^ transcriptionally CD4^+^ T cells are best described as a continuum of cell states rather than sub-clusters. This necessitated development of CSEA that will likely facilitate future single-cell transcriptomics studies with a case vs. control design. In our dataset, CSEA identified an enrichment of TFH-like signatures in MS, which we confirm independently by flow cytometry and functionally in animal models of the disease. Notably, we used a technically (i.e. active and transfer EAE) and genetically (conditional Bcl6 knock-out) considerably more rigorous approach than a previous study ^69^. We also found that TFH cells enhance B cell enrichment in the CNS in EAE and correlate with B lineage cell abundance in the CSF.

We and others^54^ thus speculate that a pathological interaction between TFH cells and B cells in the CSF may locally drive CNS autoimmune reactions. Notably, our approach is unlikely to return false positives as it is unbiased and corrected for multiple-hypothesis testing. In fact, B cell clones have long been known to,at least partially, expand in the CSF in MS^20, 70^ together with migration from the periphery^18, 40^. An importance of B cells in the disease is also supported by the expansion of plasmablasts in the CSF^41, 71^ and the efficacy of B cell-depleting therapies^21^. It will be exceptionally interesting to extend our study design to MS patients receiving B cell-depleting treatments or of MS in later disease stages (e.g. progressive). Our study provides an essential reference point for future single-cell study of human CSF and will likely facilitate understanding of diverse neurological diseases such as Parkinson’s and Alzheimer’s disease in the future.

## Supporting information

Merged Suppl Text and Figures

## Acknowledgements

We thank Claudia Kemming, Anna-Lena Börsch, Maik Höfer, Gabriele Berens, and Kirsten Weiss for technical assistance. We thank Arpita Singhal for help in developing the CSEA pipeline. G.M.z.H. was supported by grants from the Deutsche Forschungsgemeinschaft (DFG, ME4050/4-1, ME4050/8-1), from the Gemeinnützige Hertie Stiftung, from the Innovative Medical Research (IMF) program of the University Münster, and from the Ministerium für Innovation, Wissenschaft und Forschung (MIWF) des Landes Nordrhein-Westfalen. This project was funded in part by the DFG Sonderforschungsbereich Transregio 128 of the DFG (to S.G.M, A09 to H.W. and C.C.G., Z02 to H.W. and T.K.).

## Author contributions

D.S., M.H., T.L. performed experiments, M.C., M.H., C.A.X., N.Y. performed computational analyses, A.S.-M., C.G. processed CSF samples, T.K. performed histology, S.G.M., H.W. co-supervised the study, N.Y., G.M.z.H. conceived and supervised the study and wrote the manuscript. All authors critically revised the manuscript.

