## Supplementary material for "Integrated single cell analysis of blood and cerebrospinal fluid leukocytes in multiple sclerosis": Merged Suppl Text and Figures

#### Supplementary Material & Methods

*Patient recruiting and inclusion*

A total of 26 treatment-naive patients with MS or clinically isolated syndrome (CIS) receiving a lumbar puncture (LP) for diagnostic purposes, were prospectively recruited (Suppl. Table 1). The control group consisted of 22 patients diagnosed with idiopathic intracranial hypertension (IIH) (Suppl. Tab. 1). Patients were recruited in three consecutive cohorts. Cohort 1: single-cell RNA-seq of unsorted CSF cells (6 IIH vs. 6 MS patients). Cohort 2: CSF flow cytometry only (7 IIH vs. 11 MS patients), cohort 3: flow sorted $CD3^{+}CD4^{+}CXCR5^{+}$ TFH cells from CSF for bulk RNA-seq (9 IIH vs. 9 MS patients) (Suppl. Tab. 1, Suppl. Fig. 1). All patients were of Caucasian ethnicity and gave written informed consent. The study was performed in accordance with the declaration of Helsinki and approved by the local ethics committees under reference number 2015-522-f-S.

For MS patients, formal inclusion criteria were: 1) treatment naive patients with a first episode suggestive of MS (i.e. clinically isolated syndrome (CIS)) or with relapsing-remitting (RR)MS diagnosed based on MAGNIMS criteria[72], 2) patients receiving LP for diagnostic purposes and consenting to participate. Exclusion criteria for MS patients were defined as: 1) questionable diagnosis of MS by clinical signs or magnetic resonance imaging (MRI) findings, 2) secondary chronic progressive MS or primary progressive MS. IIH patients were included, if they gave informed consent. Exclusion criteria for all patients were: 1) immunologically relevant co-morbidities (e.g. rheumatologic diseases), 2) severe concomitant infectious diseases (e.g. HIV, meningitis, encephalitis), 3) pregnancy or breastfeeding, 4) younger than 18 years, 5) mental illness impairing the ability to give informed consent, 6) artificial blood contamination during the lumbar puncture resulting in >200 red blood cells / µl. Patients whose diagnostic work-up revealed a diagnosis other than MS / IIH within four weeks of clinical follow-up were retrospectively excluded (Suppl. Figure 1C). The following diagnostic tests were performed in all MS patients to exclude differential diagnoses: PCR for cytomegaly virus, Ebstein-Barr virus, Human Herpes Virus-6, Herpes simplex Virus (HSV)-1, HSV-2 and Varicella-Zoster Virus in CSF. Blood tests for anti-HAV IgM, HBsAg, anti-HBc, anti-HCV, rheuma factor, Waaler-Rose Test, anti-cyclic citrullinated peptide (CCP), antinuclear antibody (ANA), anti-double strand (ds)DNA antibodies, antineutrophil cytoplasmic antibodies (ANCA). CSF and serum were tested by the Treponema pallidum hemagglutination assay (TPHA). Borrelia burgdorferi was detected in CSF and blood by ELISA. R version 3.4.4 and RStudio 1.1.447 were used for the statistical analysis of clinical and human flow cytometry data.

*Sampling and flow cytometry analysis of cerebrospinal fluid cells and blood cells*

LPs were performed under sterile conditions using 20G Sprotte Canulae (Pajunk Medical). Up to 5 ml of CSF and 3 ml of blood were collected in addition to diagnostic material. Samples were pseudonymised at collection. CSF was quickly transported to further processing and centrifuged at 300g for 10 min. The supernatant was removed and CSF cells were resuspended in 5 ml of X-Vivo15 media (Lonza) and stored at 4°C until processing. CSF flow cytometry was performed in all donors using a Navious flow cytometer (Beckman Coulter). Blood cells were incubated in VersaLyse buffer and blood and CSF cells were stained using the following anti-human antibodies (Beckman Coulter; clone names indicated): CD3 (UCHT1); CD4 (13B8.2); CD8 (B9.11); CD14 (RMO52); CD16 (3G8); CD19 (J3-119); CD45 (J.33); CD56 (C218); CD138 (B-A38). For CSF scRNA-seq, CSF cells in media were centrifuged at 400 g for 5 min and resuspended in 40 µl of X-Vivo15 media. 5 µl of the single-cell suspension were manually counted in a Fuchs-Rosenthal chamber. The maximum of CSF cells used for input was 10,000 cells. If total available CSF cell numbers were lower than 10,000 cells, all available cells were processed. On average 5,917 cells ±

1,505 SD (control 6,167 cells ± 2,614 SD vs. MS 5,667 cells ± 1,506 SD) CSF cells were used as input per donor. For blood scRNA-seq, blood was layered on top of Lymphoprep$^{TM}$ (Stemcell) and gradient centrifugation was performed in accordance with manufacturer's instructions. After centrifugation, PBMCs enriched in the interface were taken off and washed in 10 ml of X-Vivo15 media. 5 µl of the single-cell suspension were manually counted in a Fuchs-Rosenthal chamber. The maximum of blood cells used for input was 10,000 cells. Details on cell numbers and gene capturing rates are provided in Suppl. Tab. 3.

*Generation of single-cell libraries and sequencing*
Single-cell suspensions were loaded onto the Chromium Single Cell Controller using the Chromium Single Cell 3' Library & Gel Bead Kit v2 (both from 10X Genomics) chemistry following the manufacturer's instructions. Sample processing and library preparation was performed according to manufacturer instructions using AMPure beads (Beckman Coulter). Sequencing was carried out on a local Illumina Nextseq 500 using the High-Out 75 cycle kit with a 26-8-0-57 read setup.

*Code Reproducibility*
The code for reproducing the results in this manuscript has been deposited at https://github.com/chenlingantelope/MSscRNAseq2019.git

*Preprocessing of sequencing data*
Processing of sequencing data was performed with the *cellranger* pipeline v2.0.2 (10X Genomics). Raw bcl files were de-multiplexed using *cellranger mkfastq*. Subsequent read alignments and transcript counting was done individually for each sample using *cellranger count* with standard parameters. *Cellranger aggr* was employed, to ensure that all samples had the same number of confidently mapped reads per cell. The *cellranger* computations were carried out at the High Performance Computing Facility of the Westfälische Wilhems-University (WWU) Münster.

*Single-Cell Sample Filtering*
Initial exploratory data analysis identified one MS sample and one IIH sample whose clustering did not overlap with any of the other samples (data not shown). This suggested strong batch effects. Both samples were excluded from further analysis, leaving 4 control- and 4 MS-derived samples from CSF and 5 control and 5 MS-derived samples from PBMC.
Nine barcode-level quality control (QC) metrics were computed for the unfiltered 10x Cell Ranger output: (1) number of unique molecular identifiers (UMIs), (2) number of reads, (3) mean reads per UMI, (4) standard deviation of reads per UMI, (5) percent of reads confidently mapped to the gene, (6) percent of reads mapped to the genome but not a gene, (7) percent of reads unmapped, (8) percent of UMIs corrected by the Cell Ranger pipeline, and (9) the number of cell barcodes corrected by the Cell Ranger pipeline. These metrics were used for filtering and normalization. We applied the gene and sample filtering using a scheme previously described[73]. This involved four steps:

1. Define *common genes* based on UMI counts: Genes with $n_u$ or more UMIs in at least 25% of barcodes, where $n_u$ is the upper-quartile of the non-zero elements of the UMI matrix.
2. Filter samples based on QC metrics. Remove samples with low numbers of reads, low proportions of mapped reads, or low numbers of detected common genes. The threshold for each measure is defined data-adaptively: A sample may fail any criterion if the associated metric under-performs by $z_{cut}$ standard deviations from the mean metric value or by $z_{cut}$ median absolute deviations from the median metric

value. Here we have used $z_{cut}$ = 2. This function is implemented in scone::metric_sample_filter (see below).

3. Remove barcodes from donors with fewer than 100 barcodes following sample filtering. These donors have contributed too few high-quality samples to reliably estimate donor-specific effects. Only seven cells were removed in this step.
4. Filter genes based on UMI counts: Genes with $n_u$ or more UMIs in at least $n_s$ barcodes, where $n_u$ is the upper-quartile of the non-zero elements of the sample-filtered UMI matrix. We have set $n_s$ = 5 to accommodate markers of rare populations. This sub-step ensures that included genes are detected in a sufficient number of samples after sample filtering. For the $CD4^+$-only analysis this step was applied again after the data matrix was subset to include only $CD4^+$ clusters.

*Single-Cell Harmonization*

We utilized a Bayesian variational inference model scVI [34] to infer a shared latent space of dimension 10 for all single cells from different tissue, condition and batches. Visualizations were generated using UMAP to further reduce the latent space to two dimensions. scVI is a deep generative model that learns a probabilistic representation of the transcriptional states of single-cells conditional on the sequencing batches, thus no explicit library size and batch correction is needed.

*Level 1 Clustering Analysis*

After sample filtering, we performed louvain clustering on the scVI latent space as implemented in https://github.com/taynaud/python-louvain. We first constructed a k-nearest-neighbor graph from the scVI latent space, and then used the louvain.find_partition function with the ModularityVertexPartition method to recover a total of 25 clusters. Three of these clusters correspond to CD4 T cells and were tentatively combined into a single cluster for further analysis resulting in 22 first level clusters (Suppl. Fig. 2A,B). From this, we removed one red blood cell (RBC) cluster (2,333 cells; *HBA1, HBA2, HBB*), three clusters with high doublet probability (see below) and one blood-derived cluster with low quality (mitochondrial genes, no canonical marker genes) (361 cells) for further analysis (Suppl. Fig. 2A, B)

Doublet Detection

We computed a doublet score for each single cell using the function scrub_doublets in the Scrublet package[74] with all default parameters. We then removed all clusters with greater that 20% of cells labeled as doublets (1,186, 290 and 105 cells) including one cluster of lower quality cells, one cluster expression Monocyte marker genes, and one cluster expression B cell marker genes.

Level 2 Clustering Analysis

For cells that were classified as a single cluster but two distinct clusters were visible on UMAP visualization (Monocytes, B cells and mDC cells) (Suppl. Fig. 2B, C), we performed further clustering on the scVI latent space using Spectral Clustering from the scikit-learn package SpectralClustering with number of cluster set to 2 and affinity matrix computed using k-nearest-neighbor with k=15 (Suppl. Fig. 2C). The clusters we visually identified on UMAP were confirmed to be the same as the results of Spectral Clustering. With further validation using signature genes, we included the second level clusters into the main analysis as classical monocyte, non-classical monocyte, B1, B2, mDC1 and mDC2.

T cell clustering analysis

For all CD4 T cells (excluding regulatory T cells), we performed Louvain clustering on the scVI latent space, excluding all other cells. With the same parameters as the Level 1 clustering analysis. We partitioned the CD4 T cells into a total of 12 clusters.

*VISION Analysis*

We passed raw and normalized UMI data to the VISION pipeline (https://github.com/YosefLab/VISION)[46]. Mean expression per gene symbol was calculated prior to the analysis in order to make the features relatable to general gene signatures. The goal of FastProject analysis – on which VISION is based – is to uncover biologically meaningful gene signatures that vary coherently across single-cell neighbourhoods[46]. These signatures can help assign meaning to the dominant expression differences between clusters. In addition to raw data, we passed QC, donor, status, and Seurat cluster covariates for exploratory analysis and visualization. VISION quantifies the extent to which cell signature values cluster across the cell manifold by using "consistency testing." VISION scores the extent to which neighbouring cells (similar expression profiled) are predictive of a cell's signature value using autocorrelation (Giri's C) statistics, comparing against random permutations in order to assign statistical significance with respect to a uniform null model. We also included the Seurat t-SNE as a precomputed projection. Our signature set includes:

1. Human cell cycle genes described before[9], representing sets of genes marking G1/S, S, G2/M, M, and M/G1 phases.
2. The MSigDB C7 immunological signature collection[75].
3. $T_H$ signatures compiled previously[49].
4. NetPath database signatures[76].
5. Curated T cell signatures[47].
6. Curated $T_{FH}$ [50] signature sets.
7. Curated Temra signature[48]

Housekeeping genes were referenced from the same source as the SCONE negative controls above[77].

*Comparing gene expression and cluster composition*

*Differential Composition Analysis*

For both the initial and the CD4$^+$-only clustering, we used t-test and beta-binomial generalized linear model in package aod::betabin[78] to test the difference in cluster abundances (cell counts) between MS donors and control donors. We used both methods because when cell types are rare, the estimated proportions of a cell type in each donor might be over-dispersed. The two methods show consistent results and thus we show the differential composition analysis from the beta binomial distribution comparison. For the beta-binomial regression model unless indicated in the figure legends, we set the count of the cell type of interest and the total count of cells of each donor to be the response variable and the state of the donor (MS or contro) or the tissue of origin (CSF or blood) to be the independent variable.

*Differential Expression Analysis*

We used three different tests for the discovery of differentially expressed genes between two groups of cells. First we computed Bayes Factor using the imputed counts from scVI. Bayes Factor is a generalization of the p-value and is computed as $log\frac{P(x_a \geq x_b)}{P(x_b > x_a)}$ where $x_a$ is the gene expression of the gene of interest in group $a$ and $x_a$ in group $b$. We use the generative model of scVI to obtain the batch-corrected mean of the negative binomial distribution of transcript counts. Second we used the library-size corrected UMI counts for Mann-Whitney U test. At last we followed the methods of the best performing method in a single-cell specific

DE method assessment paper[79] and we used EdgeR[33] with cellular detection rate and batch id as covariates.

*Gene Set Enrichment Analysis (GSEA)*

After deriving lists of differentially expressed (DE) genes (Suppl. Tab. 5, Suppl. Tab. 7, Suppl. Tab. 8, Suppl. Tab. 10, Suppl. Tab. 13), we applied GSEA tests[80] to all cluster specific DE gene lists DE between CSF and blood (Suppl. Tab. 6, Suppl. Tab. 11). We used the enrichr function in gseapy v0.9.12 to find overlap between the DE genes and function genesets. We used signed significance scores based on the Adjusted P-value provided by the enrichr function. Sets considered in this analysis include all MSigDB C7 signature sets and all curated T cell signature sets described previously[47] with 10 or more genes quantified in the present study; "UP" and "DN" signature subsets were tested separately.

*Cell Set Enrichment Analysis (CSEA)*

For the $CD4^+$ T cells analysis we developed a novel adaptation of the GSEA method, applying the technique to cell sets: CSEA (illustrated in Suppl. Fig. 8A). CSEA is a hypothesis testing method for simultaneously uncovering enrichments and identifying subsets of cell sets of importance. In this procedure, a collection of cells is first ordered by a transcriptional phenotype of interest (e.g., sum expression of genes in a pathway). The resulting statistical test is sensitive to cases in which only a subset of cells from one group (e.g., MS) exhibit unusually high values of the transcriptional phenotype. The input to this method is a list of *N* cells, rank-ordered by some input signal. Our analysis uses VISION signature scores, reflecting known axes of biological variation. VISION signature scores – based on FastProject signature scores[46] – are computed by first centering and scaling each normalized log expression cell profile. Following scaling, the sum of gene expression values in the negative signature subset are subtracted from the sum of gene expression values in the positive signature subset. Signatures are normalized to the total number of genes in the set. For example, a signature set that describes a dichotomy between naïve and memory T cells may be used to score individual cells, indicating that some cells have higher expression of genes characterizing the naïve state and lower expression of genes characterizing the memory state. Using the notation previously described[80] we will use $r_j$ to denote the cell j's signature score; indices have been sorted so that $r_j > r_{j+1}$ . The test involves considering all cells up to a specific position, i. A "hit" score is defined as the signature score optionally exponentiated by parameter *p* ($|r_j|^p$) for members of cell set *S*, divided by the sum over all set members in the list. A "miss" score is similarly calculated for non-members of *S*, but without weighing by signature score magnitudes.

The CSEA enrichment score (ES) is defined as the maximum of the difference between the running cumulative sum of hit scores and miss score with respect to index i. When $p$=0, the ES reduces to a one-sided KS test statistic for differential signature analysis between cell sets. When $p$=1, the cells in S are weighted by their signature score, normalized by the sum of the score over all the cells in S. We apply the same permutation scheme as described for GSEA above. For $p$>0, CSEA cannot be seen as a simple differential signature test: CSEA tests for enrichment of a cell set at the high tail of the signature score distribution, but additionally weighs the set elements according to their signature value. This reduces the effects of low-magnitude cells in *S*, whereas all cells not in *S* are treated the same no matter the magnitude of their signature score. CSEA tests if high magnitude (positive or negative) cells are enriched at a specific tail, applying permutation tests to account for the additional variability induced by the magnitude weights. The set of indices up to where the objective score reaches its maximum also holds significance – in GSEA[80] referred to as the "leading-edge" of the enrichment test. The intersection of the set *S* and the leading-edge is the *leading-edge subset*, representing an important core subset of cells driving an

enrichment. For each VISION signature, we treated the computed signature scores as cell signature scores $r_j$. The sets under consideration were the mutually exclusive sets of MS and control cells. The goal of this approach is to identify core sets of cells that drive each biological condition's enrichment for high signature values (Suppl. Fig. 8).
To screen a set of gene signatures, we computed the Vision signature score for 64 gene signatures related to CD4 T cell states, cell cycle, interlukin expression and T cell subsets. (Supplementary Table To determine the P-value of the CSEA ES score for a geneset, we shuffle the disease state labels for cells 100 times and compute the probability that the maximum ES score computed with the true labels is greater than the ES score computed with the shuffled labels. We also generated a random geneset that is matched to the original signature set in both number of genes and the average expression of each gene. This is done by finding the top 20 gene that has the closest mean expression to each gene in the original signature set, and then randomly sampling one of them. We then corrected for multiple testing using the Benjamini-Hochberg procedure to generate the corrected P-values. We filtered the result based on three criteria: the corrected P-value of the true signature set is smaller than 0.05, the corrected P-value of the control signature set is greater than 0.05, and that the leading edge is smaller than 1000. This results in 2 significant signatures, TFH and Th1. We then validated this result by computing the ES scores of 1000 matched control genesets for each signature set. We then report the P-value as the probability that the true signature set's maximum ES score being greater than the maximum ES score in the matched random genesets. We also computed the ES score for enrichment in control and found that the enrichment score is significantly larger than the control set but the leading edge is much larger than for enrichment in CSF.

*Bulk RNA-Seq of sorted TFH cells*
CSF TFH cells were sorted on a BD FACS Aria™ III cell sorter using FACS Diva™ software following manufacturer's instructions using an 85 µm nozzle and the drop delay was determined using BD Accudrop™ beads. Sorting was performed using sort precision mode "purity" for live $CD3^+CD4^+CXCR5^+$ cells. Antibodies against PD-1 (EH12.2H7) and ICOS (C398.4A) were from Biolegend. Cells were sorted directly into 1,5 ml reaction tubes containing 100 µl RNA Lysis Buffer (Zymo Research). After sorting, tubes were vortexed, briefly centrifuged and frozen at -80 °C until RNA isolation. Data were analyzed using FlowJo software v10.4.1 (Tree Star, Inc.). Samples for bulk RNA-sequencing were prepared using a modified version of the SmartSeq2 protocol[81]. Unquantified purified RNA was used as input. Reaction volumes were scaled up and the number of PCR cycles during cDNA amplification adjusted accounting for the higher number of input cells compared to the original protocol [81]. Library Preparation was done by the Next UltraII FS DNA Library Prep Kit (New England Biolabs) using 1-3 ng of cDNA as input. Sequencing for 9 MS samples and 9 IIH samples was carried out on a NextSeq500 using the High-Out 75 cycle kit (Illumina).

*Bulk expression quantification*
RNA-seq reads were aligned to the RefSeq hg38 transcriptome (GRCh38.2) using Bowtie2[82]. The resulting transcriptome alignments were processed using the RNA-Seq by Expectation Maximization (RSEM) toolkit to estimate expected counts over RefSeq transcripts [83]. Several genes were quantified multiple times due to alternative isoforms unrelated by RefSeq annotation. Before expression data normalization, the gene entry with maximum counts was selected to represent the gene in further analysis.

*Bulk RNA-seq data analysis*
Sample and gene filtering were similar to the scRNA-seq filtering method above, enforcing (> 107k reads, >10% read alignment (forced), >93.3% common genes detected; corresponding

to $z_{cut}$ = 20). A total of 5 samples were removed, leaving 13 samples. Setting $n_s$ = 1, we analysed 11,383 genes below.
For each sample, we computed transcriptome alignment and quality metrics using FastQC (Babraham Bioinformatics), Picard tools (Broad Institute), and custom scripts. Computed metrics included: (1) number of reads; (2) number of aligned reads; (3) percentage of aligned reads; (4) number of duplicate reads; (5) primer sequence contamination; (6) average insert size; (7) variance of insert size; (8) sequence complexity; (9) percentage of unique reads; (10) ribosomal read fraction; (11) coding read fraction; (12) UTR read fraction; (13) intronic read fraction; (14) intergenic read fraction; (15) mRNA read fraction; (16) median coefficient of variation of coverage; (17) mean 5' coverage bias; (18) mean 3' coverage bias; and (19) mean 5' to 3' coverage bias.
Data were normalized using SCONE. 569 positive controls were derived from MSigDB C7 entries annotated to include TFH cell types, including the most frequently included gene symbols in those entries. Negative controls for RUVg and evaluation were derived from the housekeeping gene list. Control lists were sampled down to 186 genes per list so as to match mean expression of genes in each list. The study group included two batches with 4/3 and 3/3 MS/IIH samples respectively. Biological condition was used only for evaluation. SCONE recommended TMM scaling and adjustment for 2 factors of RUVg and batch condition.
We performed PCA on the scaled log-transformed normalized data for visualization. DE between MMS and IIH donors was performed with limma-voom, using RUVg factors and batch in the model to adjust for unwanted variation. Per-gene DE significance scores were computed from log-transformed *P*-values. No single gene reached significance after correction for multiple hypothesis testing.

*Mice and EAE induction*
$CD4^{Cre}$ (ref. [84]), $2D2^{tg}$ (ref. [57]), B6.129S(FVB)-*Bcl6*$^{tm1.1Dent}$/J (named $Bcl6^{flox}$ or $Bcl6^{fl/fl}$) mice[56] were purchased from the Jackson laboratories. The $CD4^{Cre}Bcl6^{flox}$ strain was maintained by breeding the $Bcl6^{flox}$ allele to homozygosity (i.e. $Bcl6^{fl/fl}$) and breeding the Cre alleles in heterozygous to wildtype matings. Genotyping was done by routine PCR from ear punch DNA. All animal experiments were approved by the responsible state authorities (LANUV NRW) under reference number 84-02.04.2015.A319 and were performed in accordance with local regulations. Mice of both sexes (8-14 weeks old) were immunized s.c. in the flanks with an emulsion containing the myelin oligodendrocyte glycoprotein (MOG) peptide $MOG_{35–55}$ (150 µg/mouse) (GL Biochem (Shanghai) Ltd) and *M. tuberculosis* H37Ra extract (5 mg/ml, BD) in CFA (200 µl/mouse). Pertussis toxin (250 ng/mouse, Sigma) was administered intraperitoneally on days 0 and 2.
Adoptive transfer EAE induction was performed as described[85]. Briefly, naive $CD44^{low}CD62L^{high}$ $CD4^+$ T cells were FACS sorted from $2D2^{tg}$ donor mice and cultured at $2x10^6$ / ml in the presence of irradiated antigen presenting cells, soluble anti-CD3 antibody (2.5 µg/ml), IL-6 (20 ng/ml), TGF-β1 (10 ng/ml) and anti-IFNγ antibody (10 µg/ml) for 2 days. Cells were subsequently split when necessary using IL-23 (10 ng/ml) containing media for 3 additional days and then plated at $2x10^6$ / ml onto plates coated with anti-CD3 and soluble anti-CD28 (both at 2 µg/ml) in the absence of cytokines for 2 days. Cytokine production was assessed on day 5 after initial plating. Two days later, $5x10^6$ total cells were intravenously injected into C57BL/6 recipients.
Mice were monitored daily and assigned grades for clinical signs of EAE using the following scoring system: 0, healthy; 1, paralyzed tail tip; 2, paralyzed tail; 3, waddling; 4, hind legs drag on the ground; 5, butt on the ground; 6, one paralyzed hind leg; 7, both paralyzed hind legs; 8, one paralyzed front leg (criterium to stop EAE); 9, both paralyzed front legs: 10, moribund or death. Detailed refinement procedures were performed according to the

impairments of the mice. Mice with a score of >7 were euthanized. Additionally, adoptive transfer EAE recipient mice were scored with an ataxia scoring system consisting of the following criteria: ledge test, hindlimb clasping, gait and kyphosis [86]. Every criteria was rated on a scale from 0 (healthy) to 3 and all points were added up to a maximum ataxia score of 12. GraphPad Prism 5 was used for statistical analysis of all mouse-related data.

*Isolation of CNS-infiltrating mononuclear cells and lymphoid tissue characterization.*
Mice were intracardially perfused with cold PBS. The forebrain and cerebellum were dissected and spinal cords flushed out from the spinal canal with hydrostatic pressure. CNS tissue was cut into pieces and digested with collagenase D (2.5 mg/ml, Roche Diagnostics) and DNase I (0.05 mg/ml, Sigma) at 37 °C for 20 min. Mononuclear cells were isolated by passing the tissue through a 70 µm cell strainer, followed by a 70%/37% percoll gradient centrifugation. The interphase was removed, washed and re-suspended in culture medium containing 20 ng/ml PMA, 500 ng/ml ionomycin, GolgiStop, GolgiPlug (BD, each 1:1000 diluted). After 4 hours of incubation at 37 °C, cells were stained at RT for 30 min with anti-mouse antibodies (Biolegend, clones indicated): CD45 (30-11F), CD3 (17A2), CD4 (RM4-5 or GK1.5), B220 (RA3-6B2), CD19 (6D5) and live/dead staining “Zombie NIR” (BD; 1:500) in PBS. Lymph node and spleen cells were additionally stained using CD8 (53-6.7), CD11b (M1/70), CD11c (N418), Gr1 (RB6-8C5), NK1.1 (PK136) antibodies. Cells were washed and analysed using a Gallios flow cytometer (Beckman Coulter) and analysed using FlowJo V10.

*Histology*
For histology, mice were intracardially perfused with 20 ml cold PBS and fixed by perfusion with 10 ml of 4 % paraformaldehyde (PFA). Spinal cords were removed and kept in PFA for 48 hours at 4 °C. The fixed spinal cords were cut into 3 mm thick transverse segments and embedded in paraffin. To evaluate demyelination, spinal cord sections were stained with Luxol Fast Blue (LFB) and subsequently incubated with Periodic acid-Schiff (PAS). Immunohistochemistry was performed using the biotin-streptavidin peroxidase technique (K5001, Dako) in an immunostainer (AutostainerLink 48, Dako). Sections were pre-treated in a steamer (treatment solutions pH 6.0 or pH 9.0 (Dako)) before incubation with the primary antibodies against CD3 (clone CD3-12, BioRad, 1:100) or Mac3 (clone M3/84, BD, 1:100) or B220 (clone RA3-6B2, BD, 1:200). DAB was used as a chromogen. For B220/Ki67 double-immunofluorescence staining, B220 (clone RA3-6B2, BD, 1:100) and Ki67 (clone SP6, Thermo Scientific, 1:100) were used as primary antibodies; AF488- and AF594-labelled secondary antibodies (both 1:100) were used for visualization. Stained sections were analysed with a keyence microscope and pictures were taken with an Axioplot camara. ImageJ v1.48 was used to manually count infiltrated cells and measure areas.

#### Supplementary Figure Legends

*Supplementary Figure 1: Patient characteristics.*

(A) Age and gender of all control (co, n = 22) and multiple sclerosis (MS, n = 26) patients included in the study are depicted. (B) Disease characteristics of MS patients. MS patients were classified to either have (Gd+) or not have (no Gd) contrast enhancing lesions in brain or spinal cord detected by magnetic resonance imaging. Oligoclonal bands (OCB) in CSF were classified as being either undetectable (type 1), restricted to CSF (type 2), detected in serum and additionally in CSF (type 3), or not determined (?). CSF/serum indices for albumin and immunoglobulin G (IgG) were calculated. The CSF barrier function (CSF index pathology) was evaluated as being either unaffected (none), showing intrathecal IgG synthesis (Ig only), showing barrier dysfunction (barrier only), or showing both intrathecal IgG synthesis and barrier dysfunction (barrier & Ig). (C) Standard CSF parameters of all included study patients. RBC red blood cells. (D) The study recruitment scheme is depicted. 53% of control and 35% of MS samples were excluded after screening for the reasons indicated. Samples from all patients were divided into three cohorts by time of recruitment and all samples were analysed by flow cytometry. Samples in were processed for scRNAseq in cohort 1 and for bulk RNA-seq of sorted T follicular helper cells (TFH) in cohort 3. Samples in cohort 2 were only analysed by flow cytometry. (E) Clinical characteristics (age, sex) of patients *excluded* after screening and reasons for exclusion are shown. NA not applicable.

*Supplementary Figure 2: Details on the clustering approach and cell cluster identification based on scRNA-seq data.*

(A) First (1st) level clustering (see panel B) returned 22 cell cluster after tentatively merging CD4+ T cell (CD4) subclusters. Doublet, low quality, and red blood cells (RBC) were removed. Second (2nd) level clustering was performed on B cell (Bc), monocyte (mono), and mDC clusters (see panel C). Third (3rd) level clustering refers to sub-clustering of CD4+ T cells (see Fig. 2). Color codes correspond to Fig. 1B. (B) UMAP plot depicting first level clustering of all combined blood and CSF single cell transcriptomes. (C) UMAP plots showing 2nd level sub-clustering of B cells (top), monocytes (middle), mDC (bottom). (D) Feature plots of selected marker genes as in Fig. 1B. Cluster key as in Fig. 1.

*Supplementary Figure 3: Inter-individual donor heterogeneity of cell cluster abundance.*

(A) UMAP plot depicting all cells shown in Fig. 1A color-coded by tissue of origin after 2nd level clustering with blue indicating blood cells and purple indicating CSF cells. (B) Heatmap depicting cell numbers in each cell cluster in each donor sorted by tissue of origin and by disease-status. Numbers in the heatmap represent cell numbers in each cluster after 2nd level clustering. (C) Dotplot depicting selected genes differentially expressed in at least one cluster of CSF cells compared to blood. Purple dot size encodes the average expression in CSF and turquoise dot size encodes the average expression in blood. Dots are partially transparent thus overlap is dark, blue purple edge around dark blue circle indicates higher expression in CSF, while turquoise edge around dark blue circle indicates lower expression in CSF compared to blood.

*Supplementary Figure 4: Flow cytometry characterization of all CSF and blood samples.*

(A) Representative gating strategy for identifying and quantifying cell types by flow cytometry in the CSF. Population names are indicated next to the respective gates. (B) Heatmap depicting the average proportion of cells in each population in control (co, n = 22) and multiple sclerosis (MS, n = 26) patients as quantified in all samples by flow cytometry.

Heatmap color is scaled in each row to row average with color indicating higher (red) and lower (blue) than average. (C) CSF flow cytometry data are depicted as dot-boxplots if significantly (t-test statistics) different between control and MS samples. Comparisons not depicted are not significantly different. Please note that none of the analyzed flow cytometry parameters was different between co and MS in blood. (D) Flow cytometry data are depicted as dot-boxplots if significantly (t-test statistics) different between blood and CSF. * $P < 0.05$, ** $P < 0.01$, *** $P < 0.005$. *Bc* B cells, *CD4* $CD4^+$ T cells, *CD8* $CD8^+$ T cells, *dimNK* / *briNK* $CD56^{dim}$ / $CD56^{bright}$ natural killer cells, *Bc* B cells, *plasma* plasma cells, *class mono* / int mono / *nc mono* classical / intermediate / non-classical monocytes, *granulos* granulocytes.

*Supplementary Figure 5: Unlike CSF, multiple sclerosis does not affect cluster composition in blood.*
(A) UMAP plot depicting all blood cell clusters separated by disease status from control (left) and multiple sclerosis (MS, right) patients. (B) Volcano plot depicting differences of cluster abundance among all blood cells in MS samples compared with control plotting fold change (log10) against p-value (-log10) based on binomial regression modeling (Methods). Horizontal line indicates significance threshold. (C) Dotplot depicting selected genes differentially expressed in some clusters of MS cells compared to controls in CSF. Purple dot size encodes the average expression in MS and turquoise dot size encodes the average expression in controls. Dots are partially transparent thus overlap is dark, blue purple edge around dark blue circle indicates higher expression in MS, while turquoise edge around dark blue circle indicates lower expression in controls.

*Supplementary Figure 6: Late B lineage cells accumulate in the CSF in MS.*
(A) Feature plot showing the expression level of heavy chain genes IGHD/IgD, IGHM/IgM, IGHA/IgA, IGHG/IgG in the B cell clusters identified as B1/B2 in Figure 1A. Please note that IGHG summarizes IGHG1-4 genes and IGHA summarizes IGHA1-2 genes. (B) Cells expressing the respective IGH genes in the B cell clusters at maximum are highlighted. (C) Feature plots showing expression of heavy chain genes in the plasma cluster identified in Figure 1A. (D) Maximum expression of heavy chain genes in the plasma cluster.

*Supplementary Figure 7: $CD4^+$ T cells form a transcriptional continuum and show transcriptional rather than compositional changes in MS.*
(A) UMAP plot depicting sub-clustering of combined CD4+ T cells from blood and CSF as in Figure 3A with opacity of every cell reduced to 50% to improve visibility of overlaid cells (left). UMAP plot as in Figure 3A with all clusters manually pulled apart to improve visibility of overlaid cells (right). Please note that spatial localization between clusters is non informative in this plot! Outlier cells were disregarded to improve resolution. (B) UMAP plot depicting CD4+ T sub-clustered cells from CSF separated into control (left) and MS (right) cells. (C) Heatmap depicting cell numbers in each CD4+ T sub-cluster sorted by tissue of origin and by disease-status. Numbers in the heatmap represent actual cell numbers. (D) UMAP feature plots for $CD4^+$ T cells subclusters representing VISION signatures with significant VISION consistency scores ($P < 0.01$): (left) naïve vs. memory T Cell signature score[87], (right) late vs. early memory signature score[88]. (E) Volcano plot depicting differences of cluster abundance among all $CD4^+$ T cells from blood in MS samples compared with control plotting fold change (log10) against p-value (-log10) based on binomial regression modeling (Methods). (F) Heatmap showing average gene expression of selected exhaustion markers[44] in $CD4^+$ T cells combined from CSF and blood[48].

*Supplementary Figure 8: Cell set enrichment analysis (CSEA) identifies cluster-independent transcriptional changes.*

(A) Scheme of GSEA/VISION/CSEA Analysis. Available bulk expression data are used to identify gene signature sets characterizing immune cell populations (top left). These gene sets are used for either (i) gene set enrichment analysis (GSEA) of our scRNA-seq differential expression results (top middle) or (ii) single-cell VISION signature scores, input to both VISION consistency testing and cell set enrichment analysis (CSEA) testing (bottom). See Methods section for details. (B-E) Two selected CSEA analysis using TFH marker gene set (B,C) and Th1 marker gene set (D,E) in the CD4$^{+}$ T cell analysis. TFH cells ignificantly enriched in MS cells in CSF (B), but not in blood (C), while Th1 cells are enriched in both (D,E)Top row shows UMAP plots highlighting the leading edge cell sets from MS (orange) and control (green) samples. Cells depicted in grey are not part of the leading edge cell set of the respective signatures. Bottom row shows the enrichment score (y-axis) as a function of the rank of the cells by their Vision signature score (x-axis). The red line indicate the position of the leading edge. The 1D density plot shows the Vision signature score data with all MS cells in orange and control cells in green with the x-axis being the rank of the cells by their Vision signature score.

*Supplementary Figure 9: Bcl6 deficiency in T cells does not grossly affect T helper cell differentiation and secondary lymphoid organs.*

(A) Naïve CD4$^{+}$CD62L$^{high}$CD44$^{low}$ T cells were sorted from Bcl6$^{fl/fl}$ mice and CD4$^{Cre}$Bcl6$^{fl/fl}$ mice, differentiated in the presence of TGF-β1 and IL-6, or IL-12 alone, or TGF-β1 alone and analysed by intracellular cytokine staining after 4 days in culture. (B) At day 7 after active EAE induction, total live cells from spleen and draining lymph nodes (LN) from Bcl6$^{fl/fl}$ mice and CD4$^{Cre}$Bcl6$^{fl/fl}$ mice were stained for CD3, B220, CD4, CD8, CD11b, CD11c, Gr1 and NK1.1. (C) At day 28 after active EAE induction, spinal cord paraffin cross-sections were stained for LFB-Pas, Mac3 and CD3. (D) The infiltrated area (top) and number of CD3$^{+}$ cells (bottom) per spinal cord was quantified manually in a blinded fashion. (E) Cross-sections of paraffin embedded spinal cords of Bcl6$^{fl/fl}$ (left) mice and CD4$^{Cre}$Bcl6$^{fl/fl}$ (right) mice were stained for B220. Scale bars represent 100 µm in panels C and E. * $P < 0.05$, ns not significant.

*Supplementary Figure 10: Correlation between different measures of differential expression.*

(A) Matrix of Spearman rank correlation r of the different measures of differential expression of individual genes within clusters in the CSF vs. blood comparison. For example, the Bayes1 factor of all genes in the CD4 cluster is more closely correlated (i.e. yellow) with all genes in the CD8n cluster than with all genes in the Gran cluster. (B) Matrix of Spearman rank correlation r of the different measures of differential expression of individual genes within clusters in the MS vs. control comparison within CSF.

#### Supplementary Table Legends

*Supplementary Table 1: Summarized information about patients in the present study.*
Clinical characteristics (average age, sex) of all control (IIH, n=22) and multiple sclerosis (MS, n=26) patients included in the study after screening are depicted. Numbers of excluded patients for each group are also shown. All included patients were divided over three cohorts, cohort 1: CSF and blood samples used for single cell RNA-seq. (6 control vs. 6 MS), cohort 2: CSF and blood samples analysed by flow cytometry only (7 control vs. 11 MS) and cohort 3: CSF samples flow sorted for RNA-seq of $CD3^{+}CD4^{+}CXCR5^{+}$ TFH cells (9 control vs. 9 MS).

*Supplementary Table 2: Standard CSF parameters and MS disease features of patients in the present study.*
CSF parameters and MS disease features of all control (IIH, n=22) and multiple sclerosis (MS, n=26) patients included in the study are listed. All included patients were divided over three cohorts, cohort 1: CSF and blood samples used for single cell RNA-seq. (6 control vs. 6 MS), cohort 2: CSF and blood samples analysed by flow cytometry only (7 control vs. 11 MS) and cohort 3: CSF samples flow sorted for RNA-seq of $CD3^{+}CD4^{+}CXCR5^{+}$ TFH cells (9 control vs. 9 MS). All MS patients were classified if they had a relapse at CSF collection, if they had (Gd+) or not had (no Gd) contrast enhancing lesions in brain or spinal cord or if they had other MS typical characteristics observed by magnetic resonance imaging (MRI). Oligoclonal bands (OCB) in CSF were classified as being either undetectable (type 1), or restricted to CSF (type 2), or detected in serum and additionally in CSF (type 3), or not determined (?). The CSF barrier function was evaluated as being either unaffected (none), or showing intrathecal IgG synthesis (Igonly), or showing barrier dysfunction (barrier only), or showing both intrathecal IgG synthesis and barrier dysfunction (barrier & Ig) or being unclassified (unknown). Standard CSF parameters of all study patients including CSF concentrations of protein, lactate, glucose, total cells, granulocytes and red blood cells (RBC) are also depicted.

*Supplementary Table 3: Technical information of scRNA-seq results.*
Technical information on scRNA-seq results of all patients (Control, n=4 and MS, n=4) included in the study are listed. Depicted is the number of samples used for scRNA-seq (number of samples for 10x), the total number of expected cells based on counting cells included in each sample multiplied by the approximate capture rate of the 10x system (total number of expected cells), the total number of measured cells after sequencing and genome alignment (total number of measured cells), average number of measured cells per sample (average number of measured cells), average number of detected reads per cell (reads per cell) and average number of detected genes per cell (genes per cell) used for downstream analysis. The total and average number of cells measured within the $CD4^{+}$ T cell (CD4_Tc) cluster is also depicted.

*Supplementary Table 4: Differential expression analysis of all cluster for all cells.*
Genes most differentially expressed in clusters identified in the first clustering including all cells are listed. Depicted are the common Gene names (first column), the statistical values calculated by Mann-Whitney Test (stat), the Mann-Whitney test P values (pvalue), the bayes factors of upregulation in CSF/MS (bayes1), the bayes factors of upregulation in control/PBMC (bayes2), the average raw counts in CSF/MS (mean1), the average raw counts in control/PBMC (mean2), the proportion of cells expressing the genes in CSF/MS (nonz1), the proportion of cells expressing the genes in control/PBMC (nonz2), the clusters

in which this comparison is done (clusters), the log fold changes based on scVI imputed gene expression (scVI_logFC), the log fold changes based on cell size normalized counts (norm_logFC), the EdgeR P values (PValue), the EdgeR adjusted P values (FDR) and the EdgeR fold changes (logFC).

*Supplementary Table 5: Differential expression analysis of upregulated/downregulated genes in CSF vs. blood.*
Differential expression (DE) analysis was performed on CSF and blood data. Depicted are the common Gene names (first column), the statistical values calculated by Mann-Whitney Test (stat), the Mann-Whitney test P values (pvalue), the bayes factors of upregulation in CSF/MS (bayes1), the bayes factors of upregulation in control/PBMC (bayes2), the average raw counts in CSF/MS (mean1), the average raw counts in control/PBMC (mean2), the proportion of cells expressing the genes in CSF/MS (nonz1), the proportion of cells expressing the genes in control/PBMC (nonz2), the clusters in which this comparison is done (clusters), the log fold changes based on scVI imputed gene expression (scVI_logFC), the log fold changes based on cell size normalized count (norm_logFC), EdgeR fold changes (logFC), the log counts per million computed by edge R (logCPM), the statistics calculated by wilcoxon test (F), the EdgeR P values (PValue), the false discovery rates of the Mann Whitney test (fdr_wil) and the false discovery rates calculated by EdgeR (fdr_edgeR) of all upregulated (a) and downregulated (b) genes .

*Supplementary Table 6: Gene set enrichment analysis (GSEA) results for genes upregulated/downregulated in CSF vs. blood.*
Gene set enrichment analysis (GSEA) was performed on all genes (CSF vs. blood) after the first clustering (All). Depicted are the adjusted P values of the gene set enrichment analysis (Adjusted P-value), the Gene names (Gene), the GSEA terms (Term), the number of genes that significantly overlap with geneset / total number of genes in the gene set (Overlap) and the clusters they originate from (Cluster) of all upregulated (a) and downregulated (b) genes.

*Supplementary Table 7: Differential expression analysis of upregulated/downregulated genes in MS vs. IIH from blood cells.*
Differential expression (DE) analysis was performed on MS vs. IIH from blood cells. Depicted are the common Gene names (first column), the statistical values calculated by Mann-Whitney Test (stat), the Mann-Whitney test P values (pvalue), the bayes factors of upregulation in CSF/MS (bayes1), the bayes factors of upregulation in control/PBMC (bayes2), the average raw counts in CSF/MS (mean1), the average raw counts in control/PBMC (mean2), the proportion of cells expressing the genes in CSF/MS (nonz1), the proportion of cells expressing the genes in control/PBMC (nonz2), the clusters in which this comparison is done (clusters), the log fold changes based on scVI imputed gene expression (scVI_logFC), the log fold changes based on cell size normalized count (norm_logFC), EdgeR fold changes (logFC), the log counts per million computed by edge R (logCPM), the statistics calculated by wilcoxon test (F), the EdgeR P values (PValue), the false discovery rates of the Mann Whitney test (fdr_wil) and the false discovery rates calculated by EdgeR (fdr_edgeR) of all upregulated (a) and downregulated (b) genes .

*Supplementary Table 8: Differential expression analysis of upregulated/downregulated genes in MS vs. IIH from CSF cells.*
Differential expression (DE) analysis was performed on MS vs. IIH from CSF cells. Depicted are the common Gene names (first column), the statistical values calculated by Mann-Whitney Test (stat), the Mann-Whitney test P values (pvalue), the bayes factors of upregulation in CSF/MS (bayes1), the bayes factors of upregulation in control/PBMC

(bayes2), the average raw counts in CSF/MS (mean1), the average raw counts in control/PBMC (mean2), the proportion of cells expressing the genes in CSF/MS (nonz1), the proportion of cells expressing the genes in control/PBMC (nonz2), the clusters in which this comparison is done (clusters), the log fold changes based on scVI imputed gene expression (scVI_logFC), the log fold changes based on cell size normalized count (norm_logFC), EdgeR fold changes (logFC), the log counts per million computed by edge R (logCPM), the statistics calculated by wilcoxon test (F), the EdgeR P values (PValue), the false discovery rates of the Mann Whitney test (fdr_wil) and the false discovery rates calculated by EdgeR (fdr_edgeR) of all upregulated (a) and downregulated (b) genes .

*Supplementary Table 9: Differential expression analysis of genes in all cluster vs. CD4 Tc clusters.*
Differential expression (DE) analysis was performed on all clusters vs. CD4 Tc clusters including all cells. Depicted are the common Gene names (first column), the statistical values calculated by Mann-Whitney Test (stat), the Mann-Whitney test P values (pvalue), the bayes factors of upregulation in CSF/MS (bayes1), the bayes factors of upregulation in control/PBMC (bayes2), the average raw counts in CSF/MS (mean1), the average raw counts in control/PBMC (mean2), the proportion of cells expressing the genes in CSF/MS (nonz1), the proportion of cells expressing the genes in control/PBMC (nonz2), the clusters in which this comparison is done (clusters), the log fold changes based on scVI imputed gene expression (scVI_logFC), the log fold changes based on cell size normalized counts (norm_logFC), the EdgeR P values (PValue), the EdgeR adjusted P values (FDR) and the EdgeR fold changes (logFC).

*Supplementary Table 10: Differential expression analysis of upregulated/downregulated CD4 Tc cluster genes in CSF vs. blood.*
Differential expression (DE) analysis was performed on significant CD4 Tc cluster genes on CSF vs. blood. Depicted are the common Gene names (first column), the statistical values calculated by Mann-Whitney Test (stat), the Mann-Whitney test P values (pvalue), the bayes factors of upregulation in CSF/MS (bayes1), the bayes factors of upregulation in control/PBMC (bayes2), the average raw counts in CSF/MS (mean1), the average raw counts in control/PBMC (mean2), the proportion of cells expressing the genes in CSF/MS (nonz1), the proportion of cells expressing the genes in control/PBMC (nonz2), the clusters in which this comparison is done (clusters), the log fold changes based on scVI imputed gene expression (scVI_logFC), the log fold changes based on cell size normalized count (norm_logFC), EdgeR fold changes (logFC), the log counts per million computed by edge R (logCPM), the statistics calculated by wilcoxon test (F), the EdgeR P values (PValue), the false discovery rates of the Mann Whitney test (fdr_wil) and the false discovery rates calculated by EdgeR (fdr_edgeR) of all upregulated (a) and downregulated (b) genes .

*Supplementary Table 11: Gene set enrichment analysis (GSEA) results for CD4 Tc cluster genes upregulated/downregulated in CSF vs. blood.*
Gene set enrichment analysis (GSEA) was performed on significant CD4 Tc cluster genes in CSF vs. blood. Depicted are the adjusted P values of the gene set enrichment analysis (Adjusted P-value), the Gene names (Gene), the GSEA terms (Term), the number of genes that significantly overlap with geneset / total number of genes in the gene set (Overlap) and the clusters they originate from (Cluster) of all upregulated (a) and downregulated (b) genes.

*Supplementary Table 12: Cell set enrichment analysis (CSEA) results for T cell signatures.*
Cell set enrichment analysis (CSEA) was performed on all CD4+ T cells after removing residual clusters. Sheet 1 Columns in this sheet include (A) signature set (Signature), (B)

P-value for MS enrichment in CSF, (C) adjusted P-value for MS enrichment in CSF using the Benjamini-Hochberg correction, (D) control set P-values for MS enrichment in CSF, (E) control set adjusted P-value for MS enrichment in CSF using the Benjamini-Hochberg correction, (F) CSEA MS enrichment score (ES) in CSF (G) number of cells in the positive leading edge for enrichment in MS, (H-M) Same values in Blood. The P-values are computed using 100 disease state randomization. Sheet 2 shows the TFH and Th1 enrichment results including the same statistics for signature set with P-values computed only with 1000 random signature sets instead of disease state randomization.

*Supplementary Table 13: Differentially expressed genes in CSF-derived TFH cells in MS vs. control patients.*

Per-gene differential expression (DE) analysis was performed on TFH cells sorted out of the CSF from MS and control patients. The limma::topTable results for disease effect estimation are tabulated in the “DE” sheet. Columns in this sheet include i) the symbol for the gene tested (Gene Symbol), ii) $\log_e$ fold-changes (MS vs. control) in normalized expression (logFC), iii) average normalized $\log_e$-expression (AveExpr), iv) moderated t-values (t), v) statistical significance (*P*-Value), vi) Benjamini-Hochberg *Q*-value and vii) log-odds that the gene is differentially expressed. The B-value is the log-odds that the gene is differentially expressed.

### Suppl. Figure 1

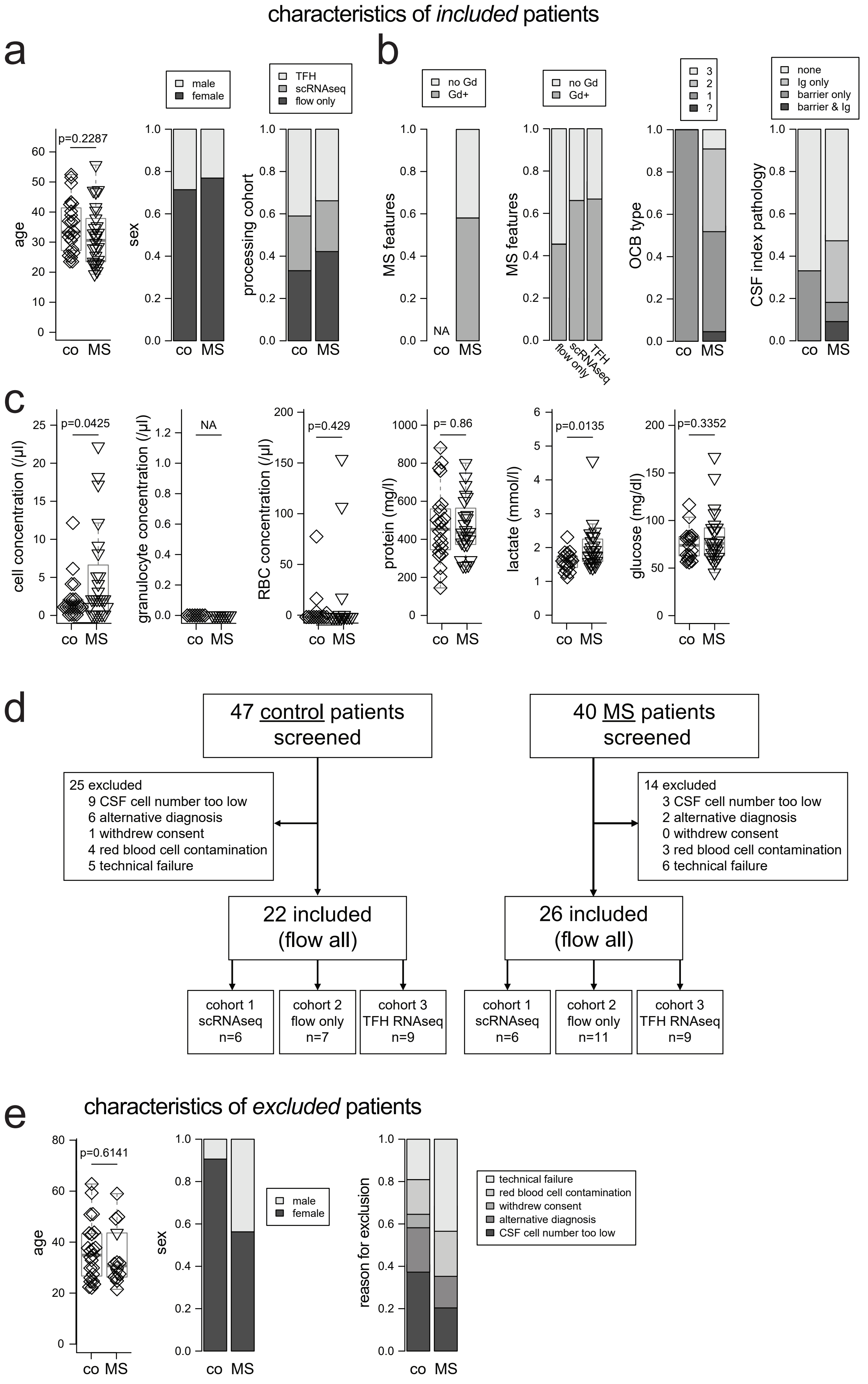

### Supplementary Figure 2

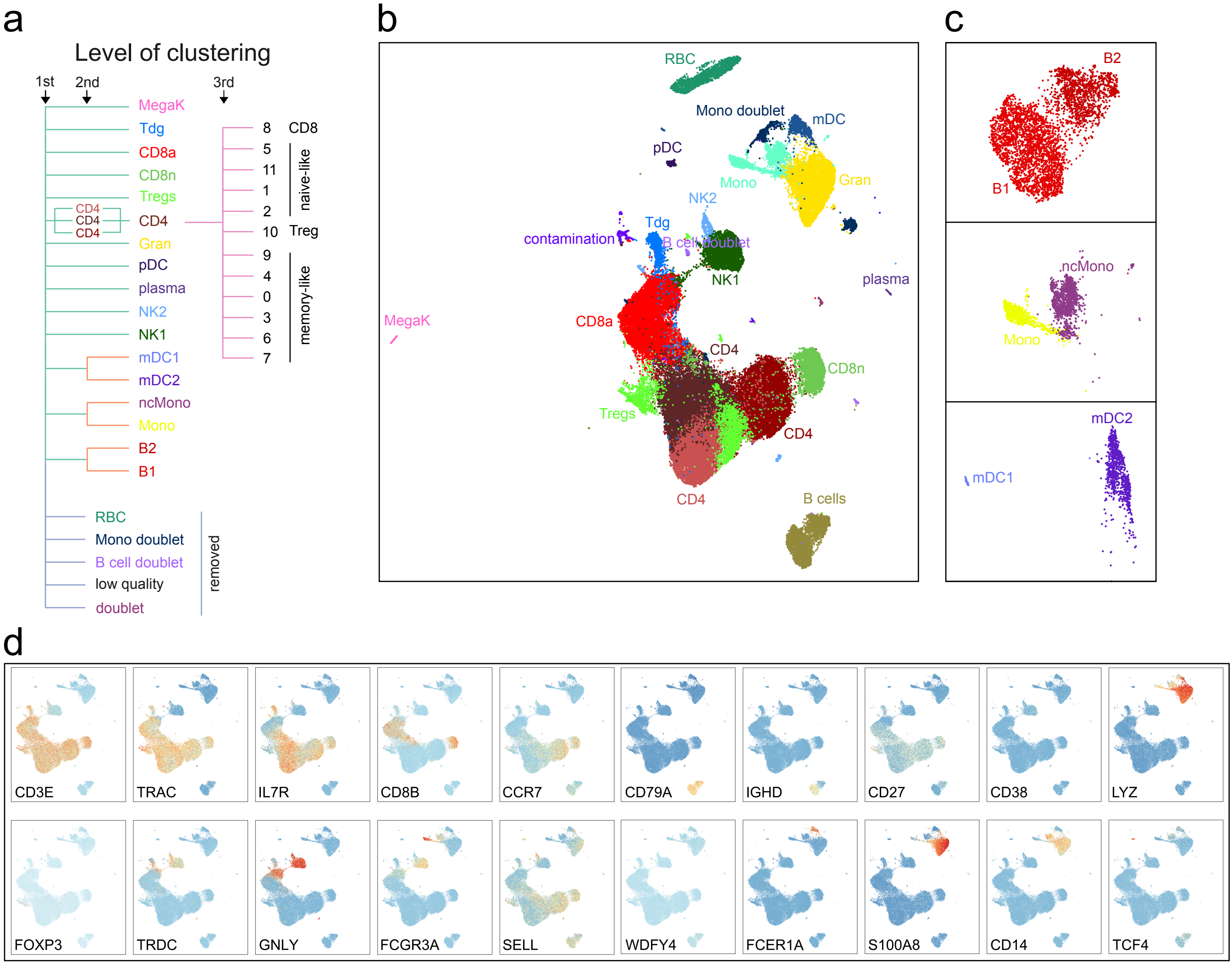

### Supplementary Figure 3

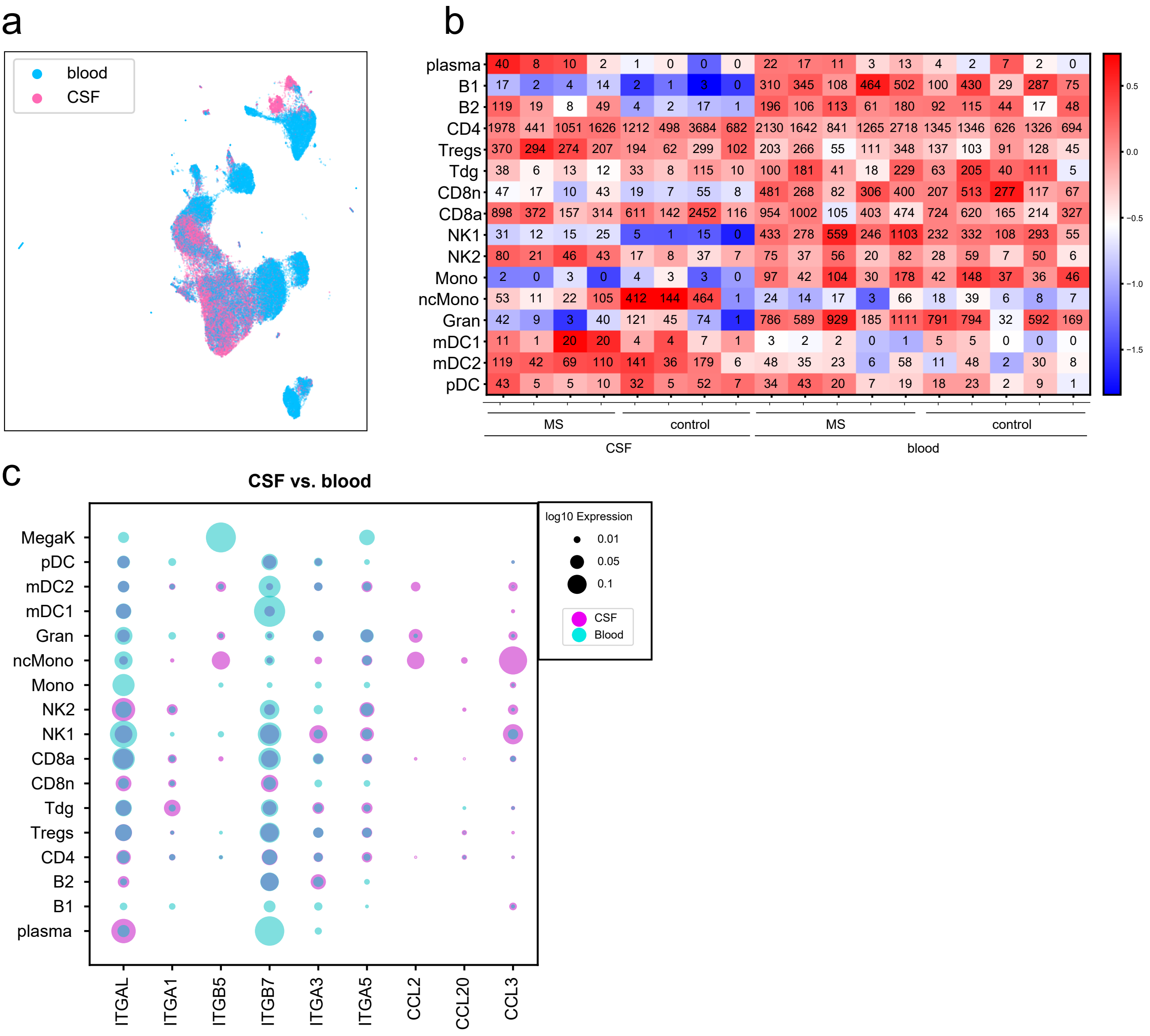

### Supplementary Figure 4

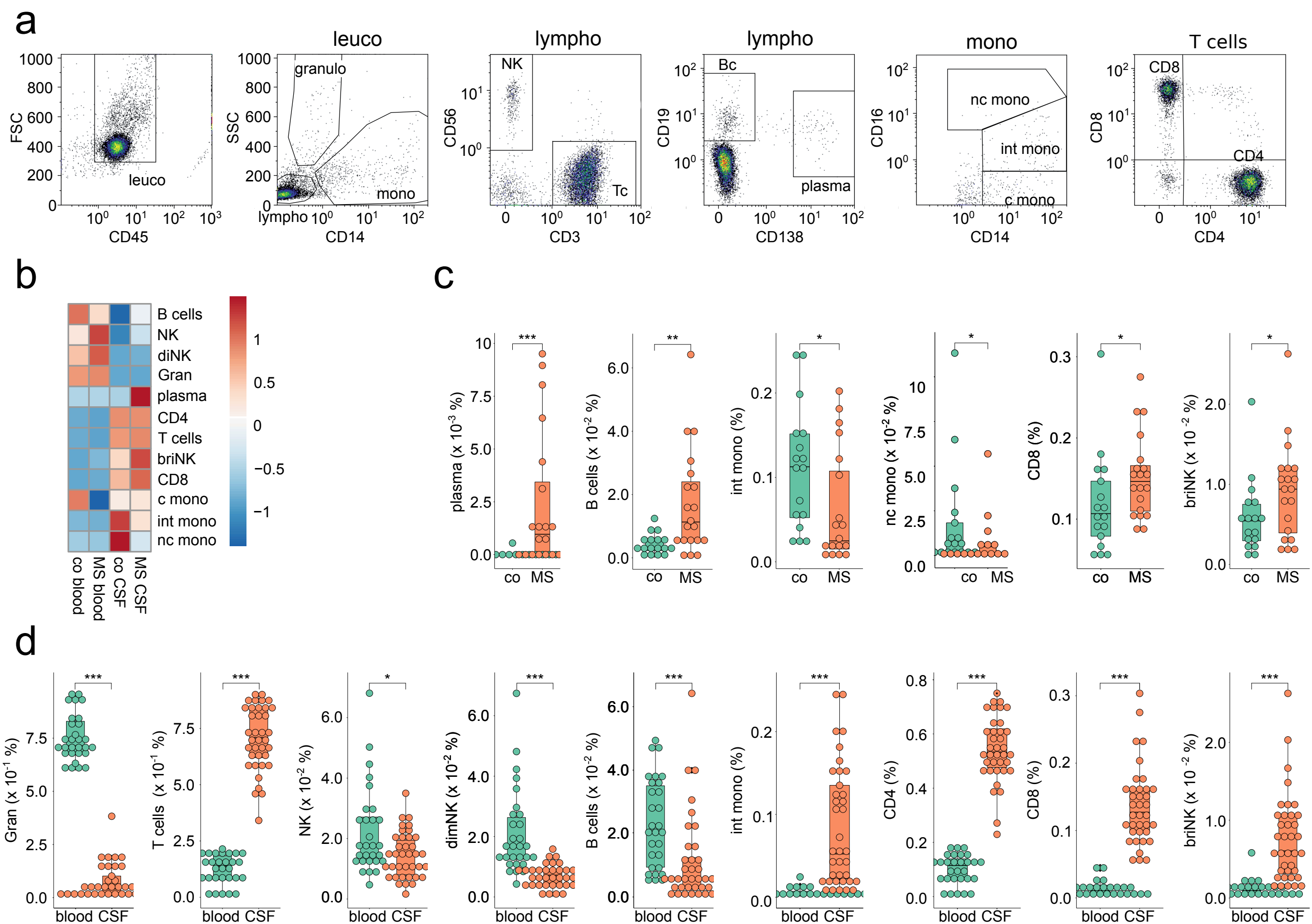

### Supplementary Figure 5

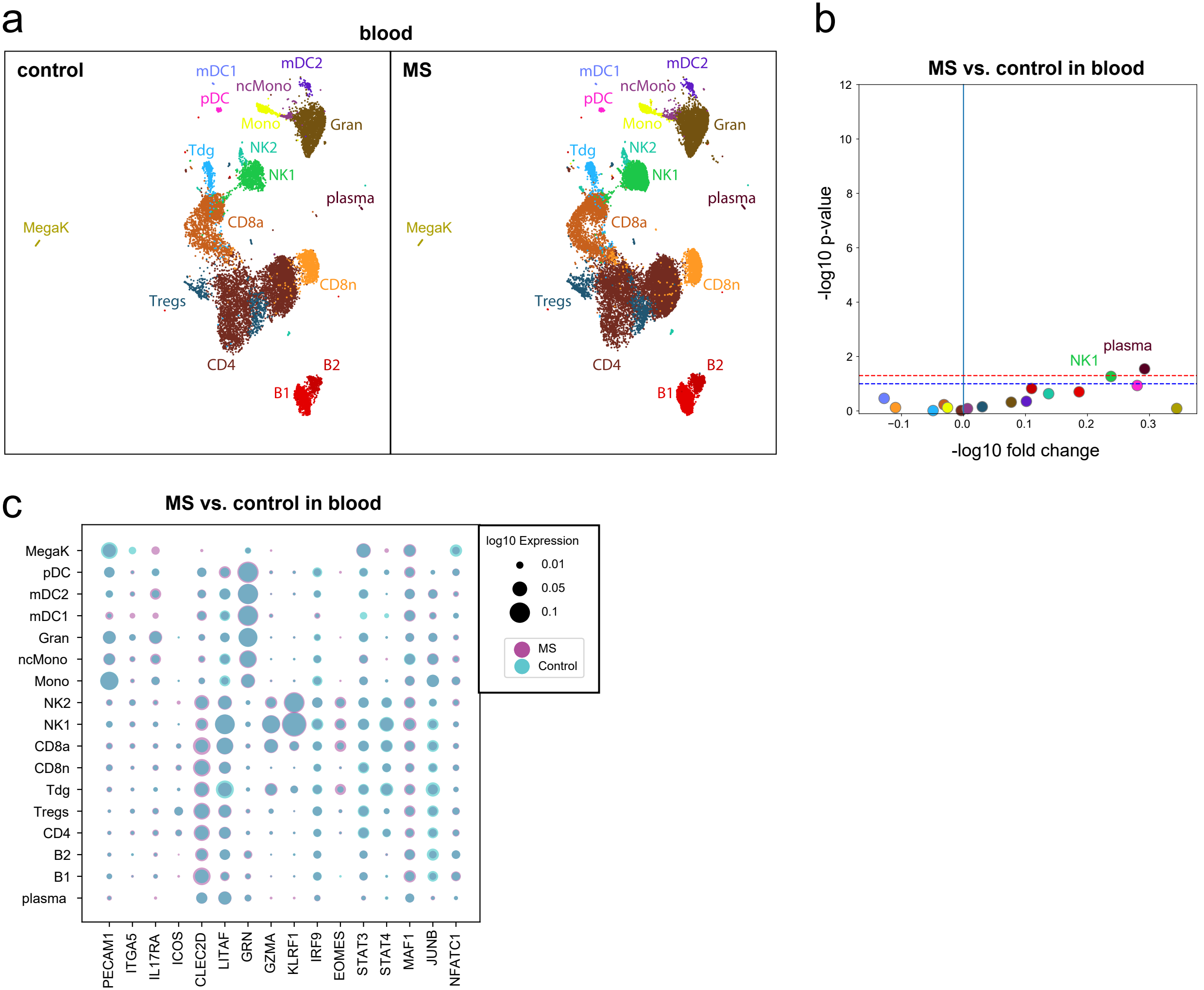

### Supplementary Figure 6

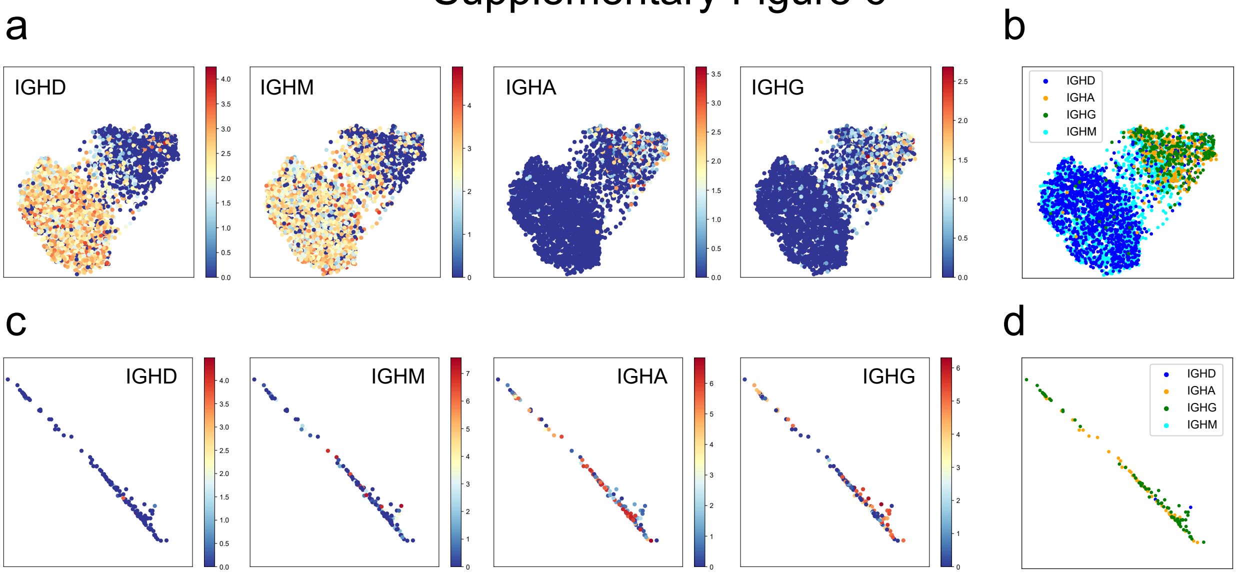

### Supplementary Figure 7

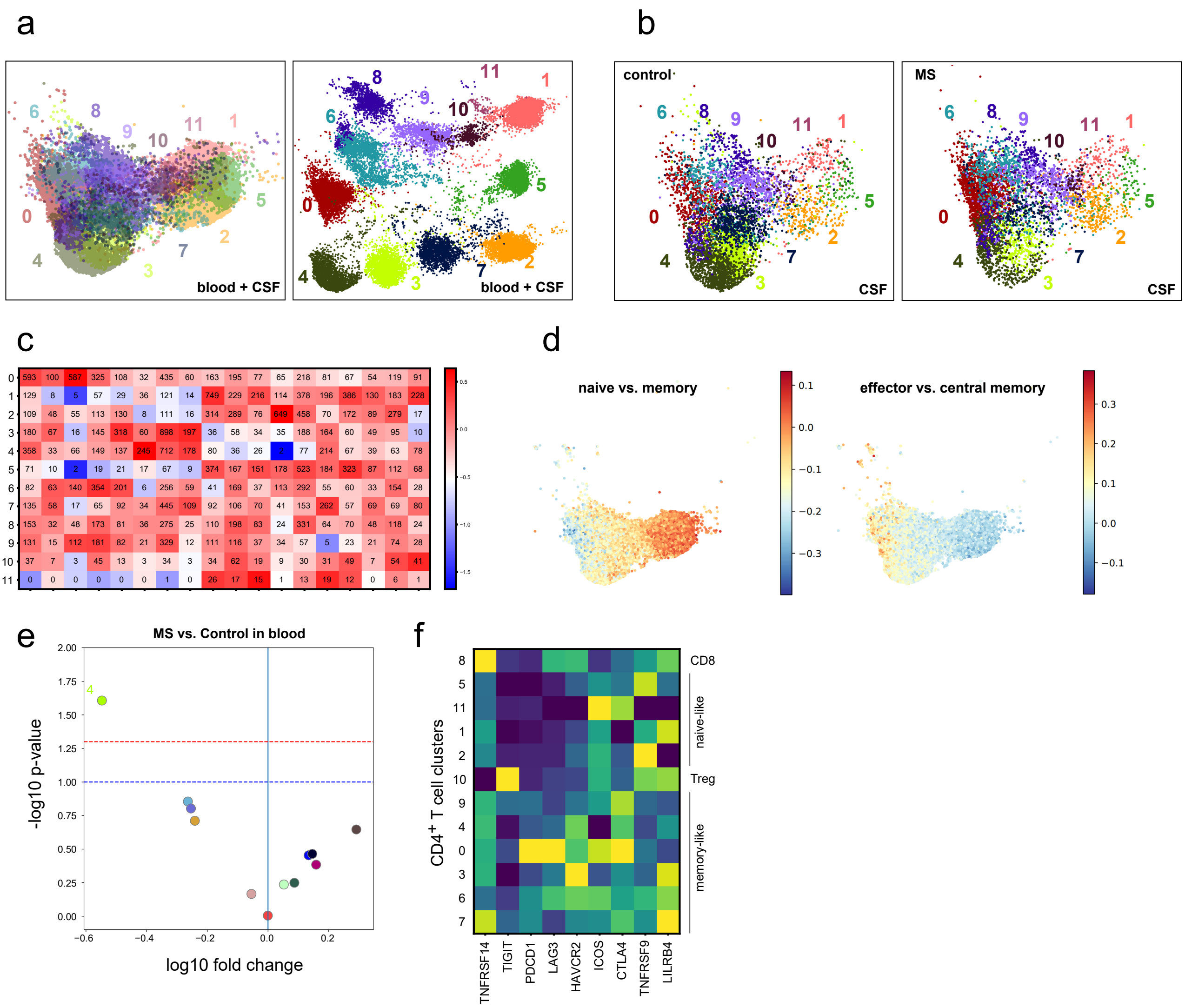

### Supplementary Figure 8

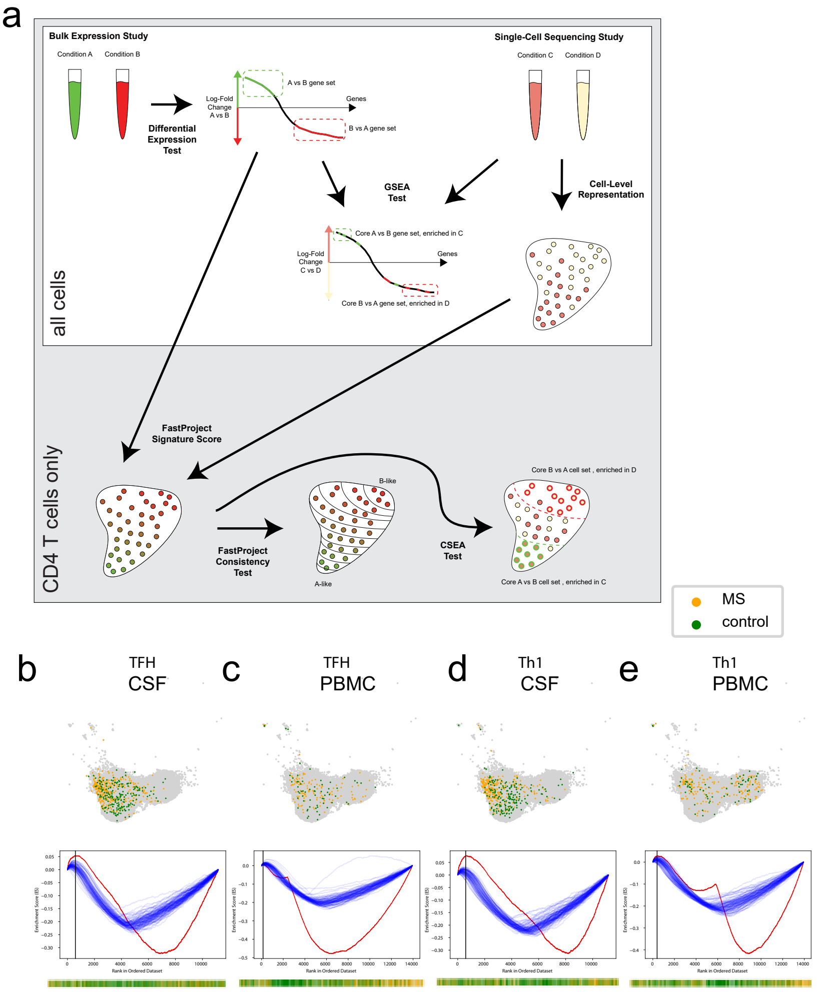

### Supplementary Figure 9

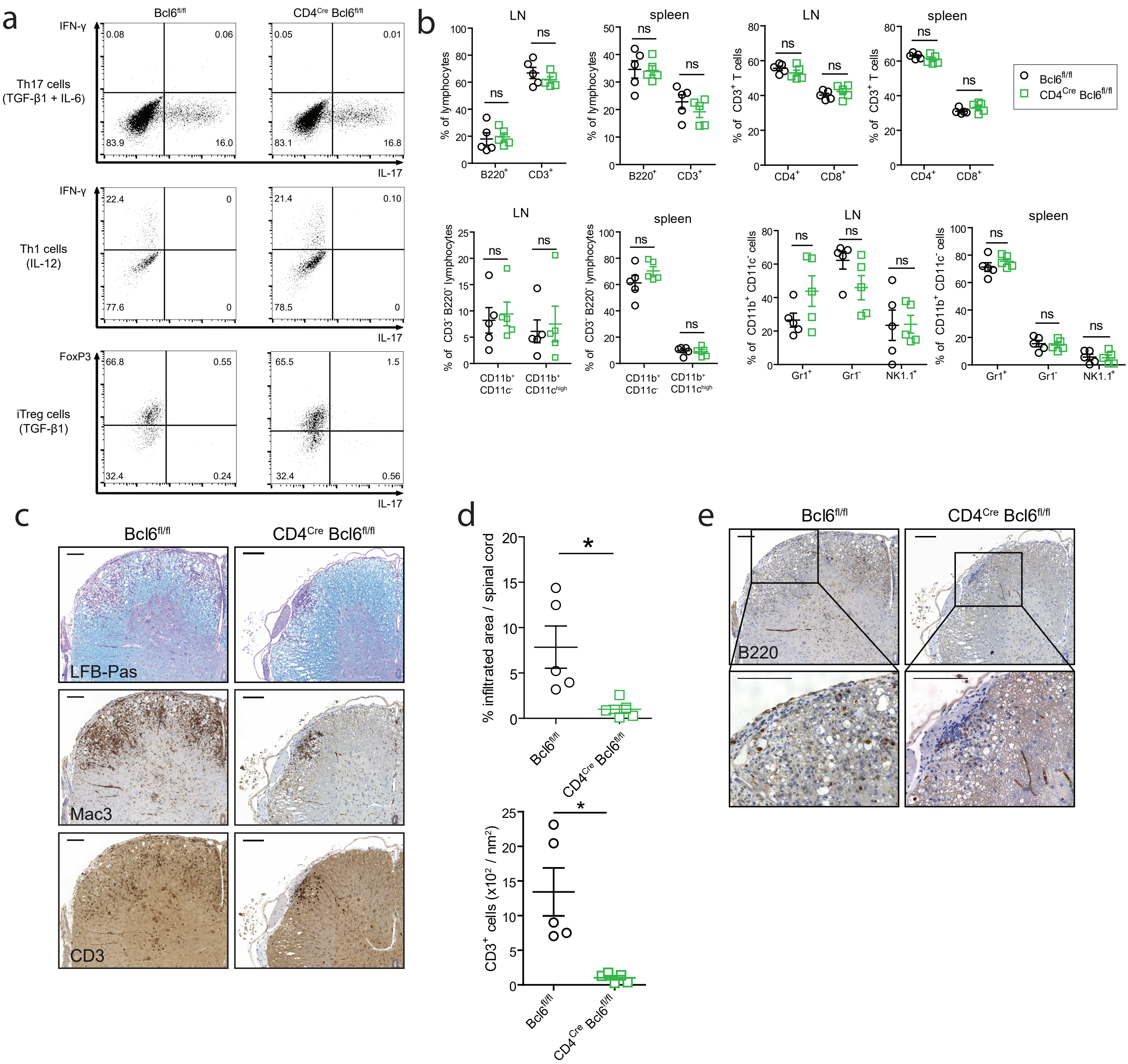

### Supplementary Figure 10

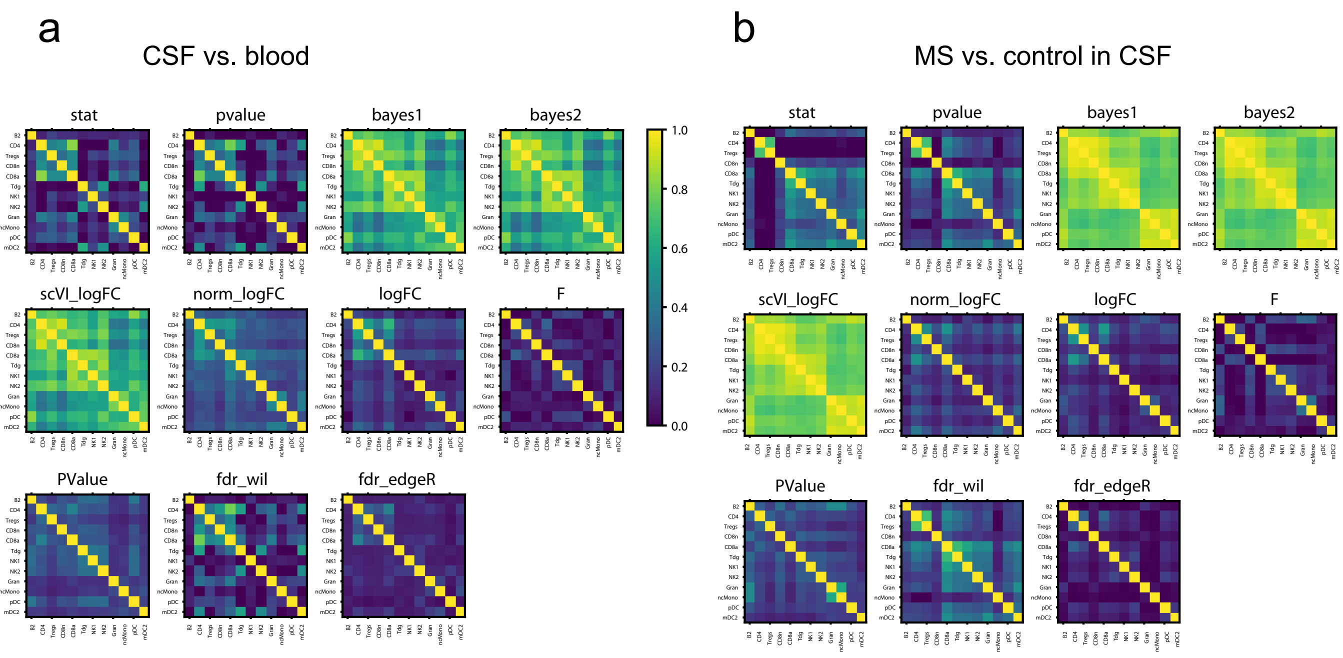
